# Glutamine metabolism modulates azole susceptibility in *Trypanosoma cruzi* amastigotes

**DOI:** 10.1101/2020.06.19.161638

**Authors:** Peter C. Dumoulin, Joshua Vollrath, Jennifer X. Wang, Barbara A. Burleigh

## Abstract

The mechanisms underlying resistance of the Chagas disease parasite, *Trypanosoma cruzi,* to current therapies are not well understood, including the potential role of metabolic heterogeneity in modulating susceptibility of intracellular amastigotes to trypanocidal compounds. We found that limiting exogenous glutamine protects actively dividing amastigotes from ergosterol biosynthesis inhibitors (azoles), independent of parasite growth rate. The antiparasitic properties of azoles are derived from inhibition of lanosterol 14α-demethylase (CYP51) in the endogenous sterol synthesis pathway. We find that carbons from ^13^C-glutamine feed into amastigote sterols and into metabolic intermediates that accumulate upon CYP51 inhibition. Consistent with a model that decreased flux through the sterol biosynthetic pathway is protective for intracellular amastigotes exposed to azoles, we find that amastigotes become re-sensitized to azoles following addition of metabolites upstream of CYP51. Our results highlight the potential role of metabolic heterogeneity in recalcitrant *T. cruzi* infection, an avenue that is currently underexplored.

## Introduction

The goal for treatment of infectious diseases caused by pathogenic bacteria or parasites is to eliminate the pathogenic microorganism from the infected host. Pathogens that persist following treatment with an antimicrobial agent may harbor genetic mutations that give rise to resistant populations. Alternatively, the pathogen may be able to achieve a dormant, non-replicative state that becomes refractory to the treatment. A third, less explored option, is the impact of metabolic and environmental heterogeneity on the efficacy of a given antimicrobial agent (Yang et al., 2017). Factors such as pathogen respiration (Lobritz et al., 2015), ATP levels (Conlon et al., 2016) and buildup of metabolic intermediates (Dumont et al., 2019) as well as environmental stressors such as the host immune response (Rowe et al., 2020) can modulate antibiotic efficacy. Recent work has shown that when the metabolic state and growth rate of microbes are disentangled, the factor that correlates with antibiotic efficacy is the microbial metabolic state (Lopatkin et al., 2019). Similarly, standard *in vitro* inhibitory activity of a candidate compound can be confounded by altered pathogen metabolism due to growth media composition (Hicks et al., 2018; Pethe et al., 2010) and conversely an understanding of these interactions can potentiate treatment (Vestergaard et al., 2017). These complex interactions are best understood in cases of bacterial pathogenesis, but recently, similar trends are apparent in eukaryotic pathogens (Dumont et al., 2019; McLean and Jacobs-Lorena, 2017; Murithi et al., 2020).

A group of single-celled protozoan pathogens with significant global disease burden exhibit metabolic and growth flexibility (Dumoulin and Burleigh, 2018; McConville et al., 2015; Saunders et al., 2010; Shah-Simpson et al., 2017) suggesting the potential for interactions with drug efficacy. In particular, kinetoplasts are a group of early branching, single celled, flagellated protists that include parasites that cause disease in humans and animals. The kinetoplastid parasite *Trypaonsoma cruzi* is the causative agent of Chagas disease and infects approximately 6 million individuals (WHO, 2015) resulting in substantial morbidity (Bern, 2015), economic burden (Lee et al., 2013) and an estimated 10,000 deaths annually (Stanaway and Roth, 2015). Parasite transmission is most common through the triatomine insect vector but can also occur orally, congenitally, by transfusion or transplantation (Bern et al., 2011; Rassi et al., 2010). Current therapies include treatment with benznidazole or nifurtimox and include undesirable characteristics such as prolonged treatment and severe adverse events (Castro and Diaz de Toranzo, 1988; Pinazo et al., 2010; Viotti et al., 2009). During the chronic stages of the disease, the elimination of parasitemia (Murcia et al., 2010) and the clinical benefit of these therapies (Morillo et al., 2015; Urbina and Docampo, 2003) are uncertain. Since the continued presence of the parasite is the main driver of disease (Jones et al., 1993; Tarleton et al., 1997; Zhang and Tarleton, 1999) a central goal for new therapies is the ability to induce sterile cure.

Azole antifungal medications that target the production of endogenous sterols were promising pre-clinical candidates (Docampo et al., 1981; Docampo and Schmuñis, 1997; Lepesheva et al., 2011; Urbina, 1997) due to the presence of erogstane-type sterols in *T. cruzi* and an already establish tolerability and safety profile in humans (Zonios and Bennett, 2008). In clinical trials it was found that monotherapy with azoles resulted in parasite suppression during treatment that was not sustained following cessation of therapy (Molina et al., 2014; Morillo et al., 2017; Torrico et al., 2018) suggesting that parasites are sensitive to therapy even in the absence of radical cure. Pharmacology and dosage do not appear to influence rebound following therapy (Khare et al., 2015a). Given the ability of intracellular *T. cruzi* amastigotes to adapt to their immediate metabolic environment (Caradonna et al., 2013; Dumoulin and Burleigh, 2018; Shah-Simpson et al., 2017) we sought to determine the extent to which this plasticity influences parasite susceptibility to ergosterol biosynthesis inhibitors. Here we show that glutamine metabolism modulates the ability of azoles to eliminate intracellular *T. cruzi* amastigotes, independent of growth rate. These protected amastigotes can be re-sensitized to azoles when supplied with exogenous sterol synthesis pathway precursors, implicating flux through the sterol synthesis pathway as a determinant of sensitivity to azoles.

## Results

### Exogenous glutamine levels modulate sensitivity of intracellular *T. cruzi* amastigotes to lanosterol-14α-demethylase inhibitors

While spontaneous emergence of latent forms of *T. cruzi* offers one possible explanation for the failure to achieve parasitological cure following drug treatment (Sánchez-Valdéz et al., 2018), the role of cellular metabolic heterogeneity in recalcitrant *T. cruzi* infection has not been explored. In previous work, we showed that proliferation of *T. cruzi* intracellular amastigotes is responsive to modifications in the exogenous growth medium and that media compositions have significant and specific interactions with small molecule inhibitors of parasite metabolism, such as the cytochrome b inhibitor GNF7686 (Dumoulin and Burleigh, 2018). To determine if changes in the metabolic environment impact the efficacy of clinically relevant compounds (Figure 1—figure supplement 1), dose-response curves were generated for benznidazole, the first line therapy for Chagas disease (Bern et al., 2007) and for ketoconazole, a potent inhibitor of trypanosome sterol synthesis (Lepesheva et al., 2011), in the presence and absence of supplemental glucose or glutamine (Figure 1A,B). We focused on these nutrients given knowledge that *T. cruzi* amastigotes are able to metabolize exogenously supplied glucose or glutamine (Shah-Simpson et al., 2017), and restriction of these exogenous sources slows amastigote replication without causing lethality (Dumoulin and Burleigh, 2018). Unlike the majority of immortalized cell lines, primary neonatal human dermal fibroblasts (NHDF/HFF) used in the present study readily withstands these conditions (Dumoulin and Burleigh, 2018). Here, we find that the dose-response observed for benznidazole is unaltered by nutrient stress (Figure 1A). Similarly, intracellular amastigotes exposed to ketoconazole in complete medium or medium lacking glucose exhibited the full range of sensitivity to ketoconazole (Figure 1B). In contrast, inhibition of intracellular amastigote growth with increasing ketoconazole levels was greatly diminished when *T. cruzi*-infected monolayers were maintained without supplemental glutamine (Figure 1B). Analogous results were obtained with other lanosterol-14α-demethylase (CYP51) inhibitors, itraconazole, posaconazole and ravuconazole (Figure 1—figure supplement 2A-C).

**Figure 1:**
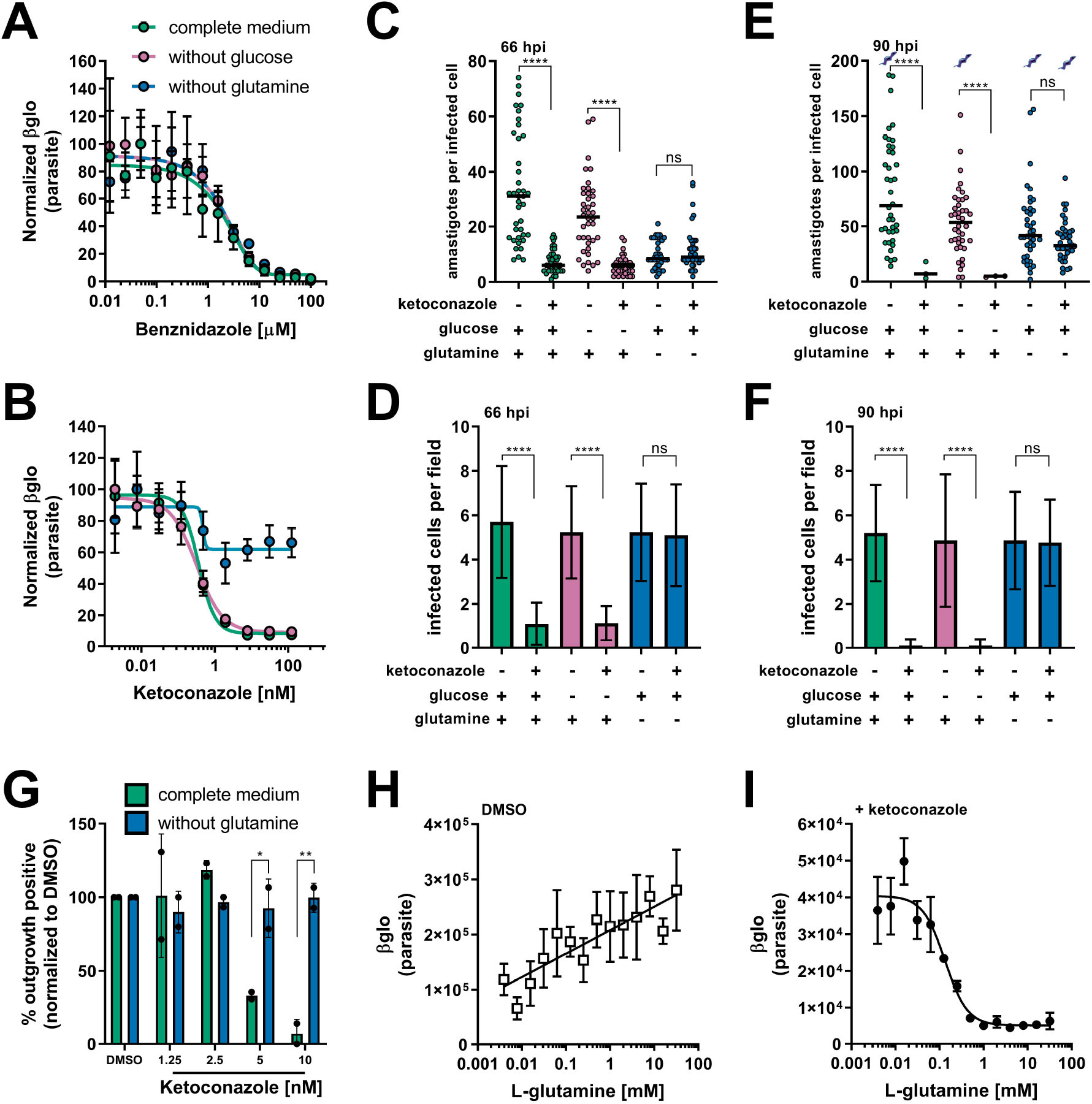
A lack of supplemental glutamine in growth medium protects intracellular *T. cruzi* amastigotes from the cytocidal effects of ketoconazole. **(A)** Dose response curves at 66 hpi of benznidazole and **(B)** ketoconazole treatment, in the indicated media compositions, normalized to the largest mean in each data set. Mean (circles) and standard deviation show (n=4). **(C)** Microscopic counts at 66 hpi of the number of amastigotes per infected host cell (n=40), medians indicated, and **(D)** the number of infected cells per field (n=20), mean and standard deviations shown. **(E)** Microscopic counts at 90 hpi of the number of amastigotes per infected host cell (n=40), medians indicated, and **(F)** the number of infected cells per field (n=20), mean and standard deviations shown. Cartoons at top of graph indicate conditions where extracellular trypomastigotes are visible in the culture supernatant. **(G)** Detection of clonal outgrowth 14 days after the indicated treatments, normalized to DMSO (vehicle) treatment. Mean and standard deviation shown, circles indicates values of two independent experiments with 28 wells used per treatment within an experiment. **(H)** Dose response curves of glutamine in the presence of DMSO or **(I)** ketoconazole (5 nM). Mean and standard deviation shown (n=3). Statistical comparisons between medians (C,E) were performed using a Kruskal-Wallis test with Dunn’s multiple comparisons test (****p<0.0001, ns=not significant). Comparisons of means (D,F) were performed using a one-way ANOVA and Bonferroni’s multiple comparisons test (****p<0.0001, ns=not significant). Comparisons of means from outgrowth (G) was performed using a two-way ANOVA with Dunnett’s multiple comparisons test (*p<0.05, **p<0.01).

To characterize the mechanism of amastigote growth suppression by ketoconazole and the protection of parasites from azoles in the absence of glutamine, we performed microscopic analysis of fixed parasite-infected fibroblast monolayers to count the number of amastigotes per infected cell (Figure 1C,E) and the proportion of infected host cells present (Figure 1D,F). Consistent with azoles acting more slowly than benznidazole (Chatelain, 2015; Dumoulin and Burleigh, 2018), no measurable effects of ketoconazole on intracellular amastigote growth following 24 hr of exposure to ketoconazole (42 hpi) were seen (Figure 1—figure supplement 3). With 48 hr of exposure to ketoconazole (5 nM, >IC_99_) (66 hpi) in complete medium or medium without supplemental glucose, there was a significant reduction in the number of intracellular amastigotes per infected cell as compared to non-treated controls (Figure 1C) and the proportion of infected cells (Figure 1D) indicative of parasite death. In contrast, intracellular amastigotes survived ketoconazole treatment under conditions of glutamine restriction (Figure 1C,D; 66 hpi) and continued to replicate as evidenced by the greater number of amastigotes per infected cell at 90 hpi (Figure 1E) without a reduction in the proportion of infected cells (Figure 1F). Furthermore, the detection of extracellular trypomastigotes in the supernatants of untreated cultures and in those treated with ketoconazole in the absence of glutamine at 90 hpi (Figure 1E; symbols), demonstrates that these ketoconazole-treated amastigotes complete the intracellular cycle in mammalian host cells to produce trypomastigotes.

To evaluate the longer-term impact of ketoconazole exposure on intracellular *T. cruz*i amastigotes cultured in the absence of supplemental glutamine, a clonal outgrowth assay was utilized to quantitatively measure parasite rebound following treatment (Dumoulin and Burleigh, 2020, 2018). As detection of outgrowth (>14 days) requires surviving parasites to successfully complete several lytic cycles, this approach distinguishes cytostatic from cidal effects of a test compound.

Exposure of intracellular amastigotes to increasing concentrations of ketoconazole in complete medium (from 18 hpi - 66 hpi) results in a proportional decrease in clonal outgrowth (Figure 1G), consistent with irreversible cytotoxicity incurred by exposure to ketoconazole (Goad et al., 1989). In contrast, no evidence of killing was seen when supplemental glutamine was restricted during the period of ketoconazole exposure as clonal outgrowth was comparable to vehicle-treated controls under these conditions (Figure 1G). Extending the ketoconazole exposure time to 72 hrs does not alter the outcome (Figure 1—figure supplement 4). These results confirm that intracellular amastigotes are protected from the lethal effects of ketoconazole upon restriction of supplemental glutamine and further demonstrate protection at the population level as opposed to the selection of a surviving amastigote sub-population that is intrinsically refractory to the drug.

### Glutamine supplementation sensitizes intracellular *T. cruzi* amastigotes to ketoconazole in a dose-dependent manner

Intracellular *T. cruzi* amastigotes succumb to the toxic effects of azoles when glutamine (2 mM) is present in the *in vitro* growth medium (Figure 1A-G). Given that standard glutamine concentrations in culture medium (1-2 mM) are significantly higher than the normal human plasma physiologic range (Cruzat et al., 2018), supplemental glutamine was titrated to determine the range in which intracellular amastigotes became sensitized to ketoconazole. In the absence of drug, the intracellular parasite load increases linearly with the addition of glutamine (Figure 1H), but in the presence of a fixed concentration of ketoconazole (5 nM, >IC99), supplemental glutamine decreased amastigote growth in a dose-dependent manner (IC_50_ of 133.4 μM for glutamine; Figure 1I). Addition of amino acids (proline/histidine), not present in the base medium, failed to sensitize amastigotes to ketoconazole and impact parasite growth (Figure 1—figure supplement 5) suggesting that glutamine metabolism in the parasite, host cell or both, plays a key role in the susceptibility of intracellular *T. cruzi* amastigotes to azole drugs.

Since intracellular *T. cruzi* amastigote growth is markedly reduced under conditions of glutamine restriction *in vitro*, we cannot rule out the possibility that slowed growth itself might protect parasites from lethality associated with blocking sterol biosynthesis with azoles. To assess whether slowed growth is an underlying factor in the protection of intracellular amastigotes from azole-mediated death, we exploited a small molecule inhibitor of parasite cytochrome b, GNF7686 (Khare et al., 2015b), which acts cytostatically, to slow amastigote replication in a dose-dependent manner (Dumoulin and Burleigh, 2018). At 66 hpi both the amastigotes per infected cell and proportion of infected cells is comparable between protection from glutamine withdrawal and GNF7686 treatment (Figure 2A,B). However, by 90 hpi when GNF6786 was applied to slow intracellular amastigote growth in complete medium, the parasites succumbed to ketoconazole treatment (Figure 2C,D *glut +, keto +, GNF +*) as did the more rapidly proliferating parasites in the absence of GNF7686 (Figure 2C,D, *glut +, keto +, GNF −*). Parasites cultured in the absence of supplemental glutamine were protected from the cytotoxic consequences of ketoconazole in GNF7686-treated (Figure 2C,D, *glut −, keto +, GNF +*) and untreated conditions (Figure 2C,D, *glut −, keto +, GNF −*). Together with the observation that restriction of supplemental glucose, another amastigote growth-limiting condition (Dumoulin and Burleigh, 2018), fails to protect intracellular parasites from ketoconazole (Figure 1B-F), these findings support the conclusion that growth inhibition of *T. cruzi* alone does not account for azole-refractory infection under conditions of glutamine restriction *in vitro.* Furthermore, generation of reactive oxygen species (ROS) due to glutamine deprivation (Matés et al., 2002) or cytochrome b inhibition (Dröse and Brandt, 2008; Fridovich, 1978) does not play a role in protection of intracellular *T. cruzi* amastigotes from azole-mediated cytotoxicity given that antioxidant supplementation does not alter the susceptibility of amastigotes to azoles under any of the conditions tested (Figure 2—figure supplement 1A,B). Similar outcomes were achieved when experiments were conducted under normoxic (~20% O_2_) or hypoxic (1.3% O_2_) conditions (Figure 2—figure supplement 1C-E). Thus, our results point to dysregulated glutamine metabolism, rather than slowed parasite growth, in the protection of intracellular *T. cruzi* amastigotes from death following exposure to azoles.

**Figure 2:**
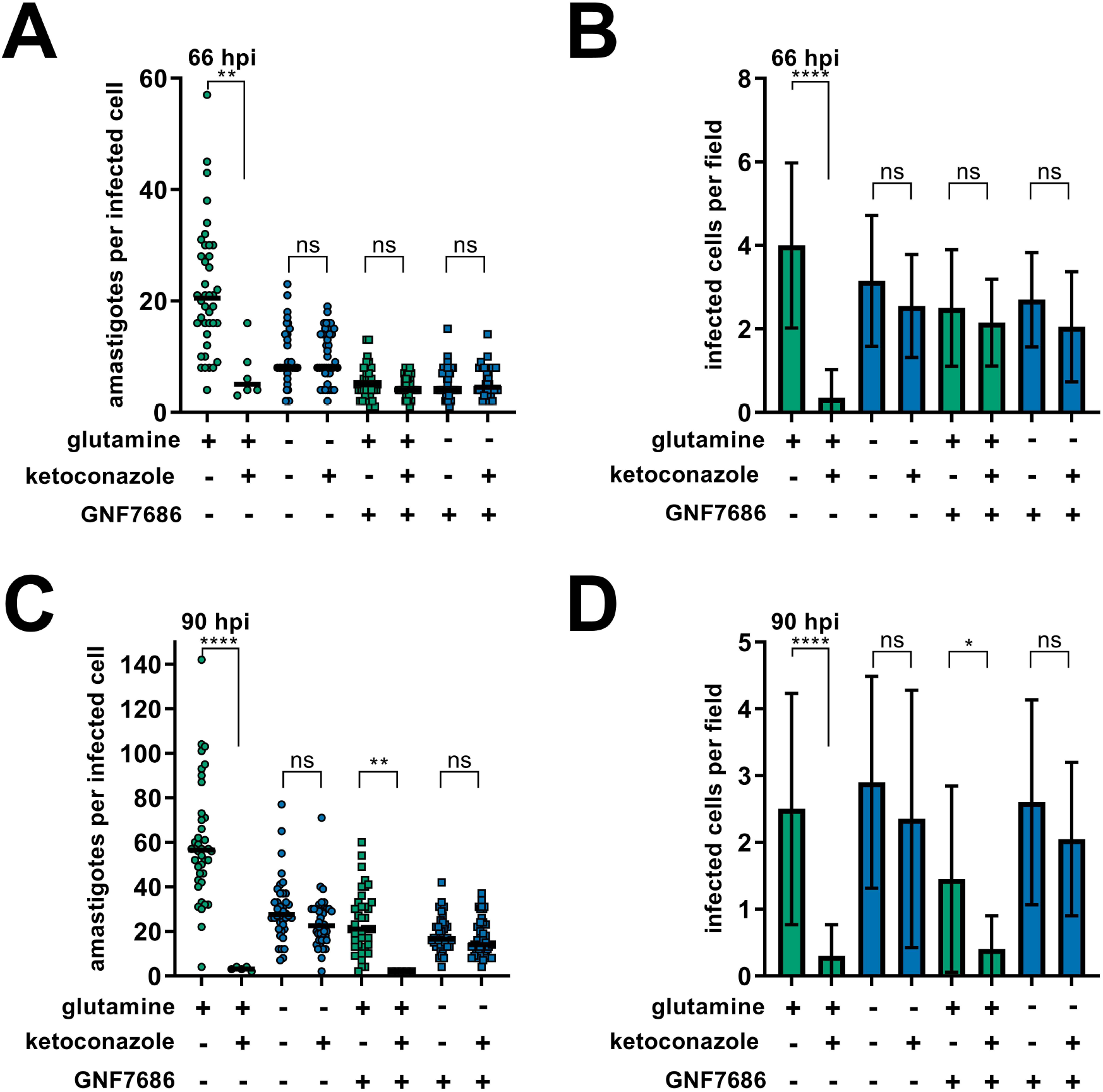
Slowed amastigote growth delays but does not prevent the cidal effects of ketoconazole. **(A)** Microscopic counts of amastigotes per host cell (n=40) and **(B)** proportion of infected cells (n=20) at 66 hpi following treatment at 18 hpi with ketoconazole (5 nM) and/or GNF7686 (150 nM) under the indicated conditions. **(C)** Microscopic counts of amastigotes per host cell (n=40) and **(D)** proportion of infected cells (n=20) at 90 hpi following treatment at 18 hpi with ketoconazole (5 nM) and/or GNF7686 (150 nM) under the indicated conditions. Statistical comparisons between medians (A,C) were performed using a Kruskal-Wallis test with Dunn’s multiple comparisons test (****p<0.0001, **p<0.01, ns=not significant). Comparisons of means (B,D) were performed using a one-way ANOVA and Bonferroni’s multiple comparisons test (****p<0.0001, *p<0.05, ns=not significant).

### Glutamine-derived carbons are incorporated into amastigote sterols

*T. cruzi* amastigotes replicate in the cytosol of their mammalian host cells and little is known regarding the interchange of metabolic precursors and metabolites between the two cells. However, the fact that the growth rate of these intracellular parasites responds to the availability of glutamine in the culture medium (Figure 1; and (Dumoulin and Burleigh, 2018)) and that isolated amastigotes are able to use glutamine as a substrate to fuel oxidative phosphorylation (Shah-Simpson et al., 2017) highlight the potential for intracellular *T. cruzi* amastigotes to take up glutamine. In this scenario, the availability of exogenous glutamine could influence the production of endogenous parasite sterols or pathway intermediates in the intracellular parasites. By performing metabolic labeling of *T. cruzi*-infected fibroblast monolayers, and isolation prior to amastigote death (Figure 3—figure supplement 1) a metabolic link between exogenously supplied glutamine and endogenous sterol synthesis in intracellular amastigotes was established (Figure 3). Carbons from exogenously supplied L-[^14^C(U)]-glutamine were found to be incorporated into sterols extracted from isolated intracellular amastigotes following separation by thin layer chromatography (TLC) (Figure 3A). A single strongly labeled sterol species, or a collection of co-migrating amastigote-derived species, present in the untreated control, were absent in amastigotes treated with ketoconazole (Figure 3A) providing evidence that sterol synthesis species downstream of CYP51 incorporate carbons from exogenous glutamine. This species, or collection of species, co-migrated with the ergosterol standard. However, *T. cruzi* amastigotes reportedly do not generate canonical ergosterol as the final species from endogenous sterol synthesis (Gunatilleke et al., 2012; Liendo et al., 1999; Ottilie et al., 2017).

**Figure 3:**
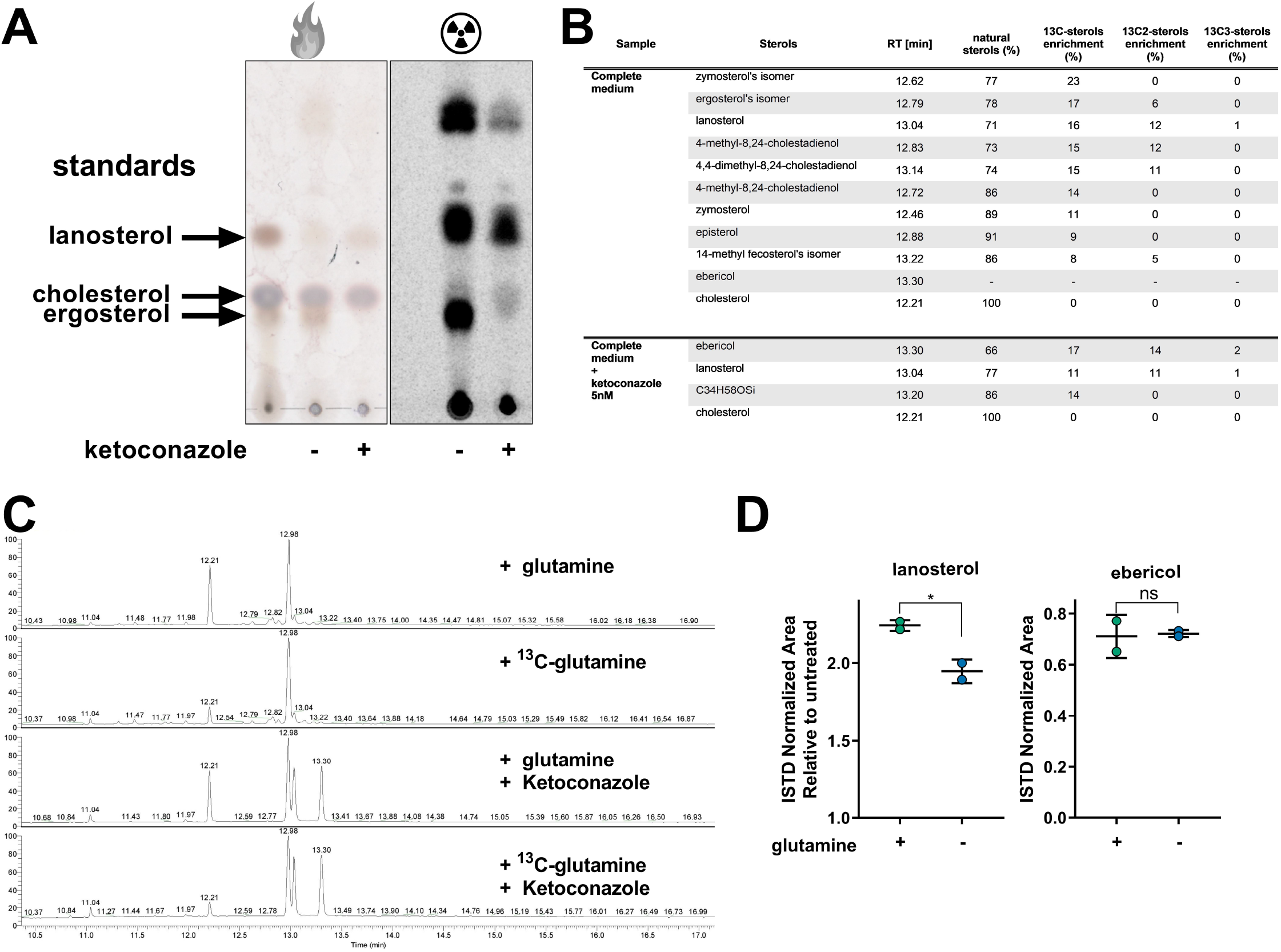
Glutamine derived carbons are incorporated into amastigote sterols and influence the buildup of lanosterol. **(A)** Thin layer chromatography (TLC) of sterols extracted from amastigotes isolated at 52 hpi with or without ketoconazole (5 nM) treatment. Cultures we spiked with 25 uCi universally labeled ^14^C-glutamine per condition at 18 hpi. Charred TLC plate for total carbons (left) and developed phosphorimager screen (right) for radioactivity with non-radioactive chemical standards are shown. **(B)** Table of detectable endogenous isolated amastigote sterol species and the percentage of ^13^C-glutamine incorporation. **(C)** Chromatogram from GC-MS detection of samples in panel B. Host cell derived cholesterol is seen at retention time 12.21, ebericol at 13.30, lanosterol at 13.04 and the internal standard at 12.98. **(D)** Quantification, using an internal standard, of lanosterol and ebericol in isolated amastigotes (52 hpi) following treatment with ketoconazole (5 nM) at 18 hpi with or without glutamine (2 mM). Mean and standard deviation shown of independent treatments, infections and amastigote isolations (n=2). Statistical comparisons are made using a Student’s t-test (*p<0.05, ns=not significant).

To identify individual sterols and the species that incorporate carbons from exogenous glutamine, we performed GC-MS following metabolic labeling of *T. cruzi* infected cultures with universally labeled ^13^C-glutamine. As expected, carbons from exogenous ^13^C-glutamine were readily incorporated into species downstream of CYP51 (Figure 3B). In the presence of ketoconazole, when CYP51 is inhibited, these downstream species were absent, and ^13^C was incorporated into lanosterol and ebericol (Figure 3B,C), the two species immediately upstream of CYP51 in the sterol synthesis pathway (Gunatilleke et al., 2012; Ottilie et al., 2017). Unlike endogenous sterols and synthesis intermediates derived from amastigote metabolism, we did not identify any incorporation of ^13^C-glutamine into host derived, but amastigote-associated, cholesterol (Figure 3B,C).

Therefore, the incorporation of carbons from exogenous glutamine into parasite sterol synthesis and in the buildup of synthesis intermediates due to CYP51 inhibition suggests that levels of 14-methylated sterol intermediates are a potential modulator of amastigote susceptibility to azoles. Optimization of the GC-MS/ISTD sterol quantification of ketoconazole treated amastigotes found that unlike lanosterol and ebericol, cholesterol levels were variable between biological replicates (coefficient of variation > 0.3; Figure 3—figure supplement 2) and could not reliably be used for comparisons between conditions. However, the variation in lanosterol and ebericol recovered from isolated amastigotes was within an acceptable range (CoV <0.3; Figure 3—figure supplement 2) and are therefore useful for comparison. In all conditions tested, ketoconazole treatment resulted in an increase in lanosterol in relation to untreated controls and the generation of ebericol (Figure 3D). We found that in the presence of ketoconazole amastigotes grown in the absence of glutamine have a reduction in free lanosterol but not ebericol (Figure 3D) suggesting that the amount of free 14-methylated intermediates and potentially the rate of generation of these intermediates are altered in parasites experiencing glutamine restriction.

### Supplementation with FPP/Farnesol is sufficient to re-sensitize amastigotes to ketoconazole in the absence of glutamine

The generation of 14-methylated sterol precursors has been implicated in the detrimental phenotypes associated with inactivation of CYP51 in other kinetoplastid protozoan parasites and yeast (Goad et al., 1989; Kelly et al., 1995; Mukherjee et al., 2019). If flux and/or generation of these methylated intermediate species modulates the sensitivity of *T. cruzi* amastigotes to azoles, we reasoned that provision of metabolites downstream of glutamine but upstream of CYP51 could re-sensitize the parasites to ketoconazole under conditions of glutamine restriction (Figure 4A). Addition of a cell-permeable form of α-KG, dimethyl α-ketoglutarate (di-α-KG), resulted in a significant reduction of intracellular amastigote growth following treatment with ketoconazole in the absence of glutamine (Figure 4B,C), while addition of di-α-KG had no effect on intracellular amastigote growth in glutamine-depleted or replete medium in the absence of ketoconazole (Figure 4B,C). This result supports the idea that bypassing the loss of exogenous glutamine by supplementing the medium can re-sensitize amastigotes to ketoconazole. However, given the potential for conversion of α-KG to glutamate and then to glutamine, it is difficult to draw definitive conclusion with regard to metabolite flux. We therefore examined the possibility that delivery of an upstream sterol precursor would also sensitize amastigotes to ketoconazole in the absence of glutamine without the potential for conversion to glutamine. In *T. cruzi,* isoprenoid precursors can enter the endogenous sterol synthesis pathway (Cosentino and Agüero, 2014) and in other systems, exogenous supplementation of isoprenoid precursors is sufficient to chemically rescue blockage of essential metabolic function (Yeh and DeRisi, 2011). In the absence of ketoconazole, addition of farnesyl pyrophosphate (FPP) (Figure 4D,E and —figure supplement 1) or farnesol (Figure 4F,G and —figure supplement 1) to the culture medium had no effect on amastigote growth. In contrast, intracellular amastigotes failed to survive ketoconazole treatment in glutamine-free medium when FPP (Figure 4D,E and —figure supplement 1) or farnesol (Figure 4F,G and —figure supplement 1) were present. These findings suggest that changes to flux through the sterol synthesis pathway is a causative factor for both protection from and sensitivity to azoles.

**Figure 4:**
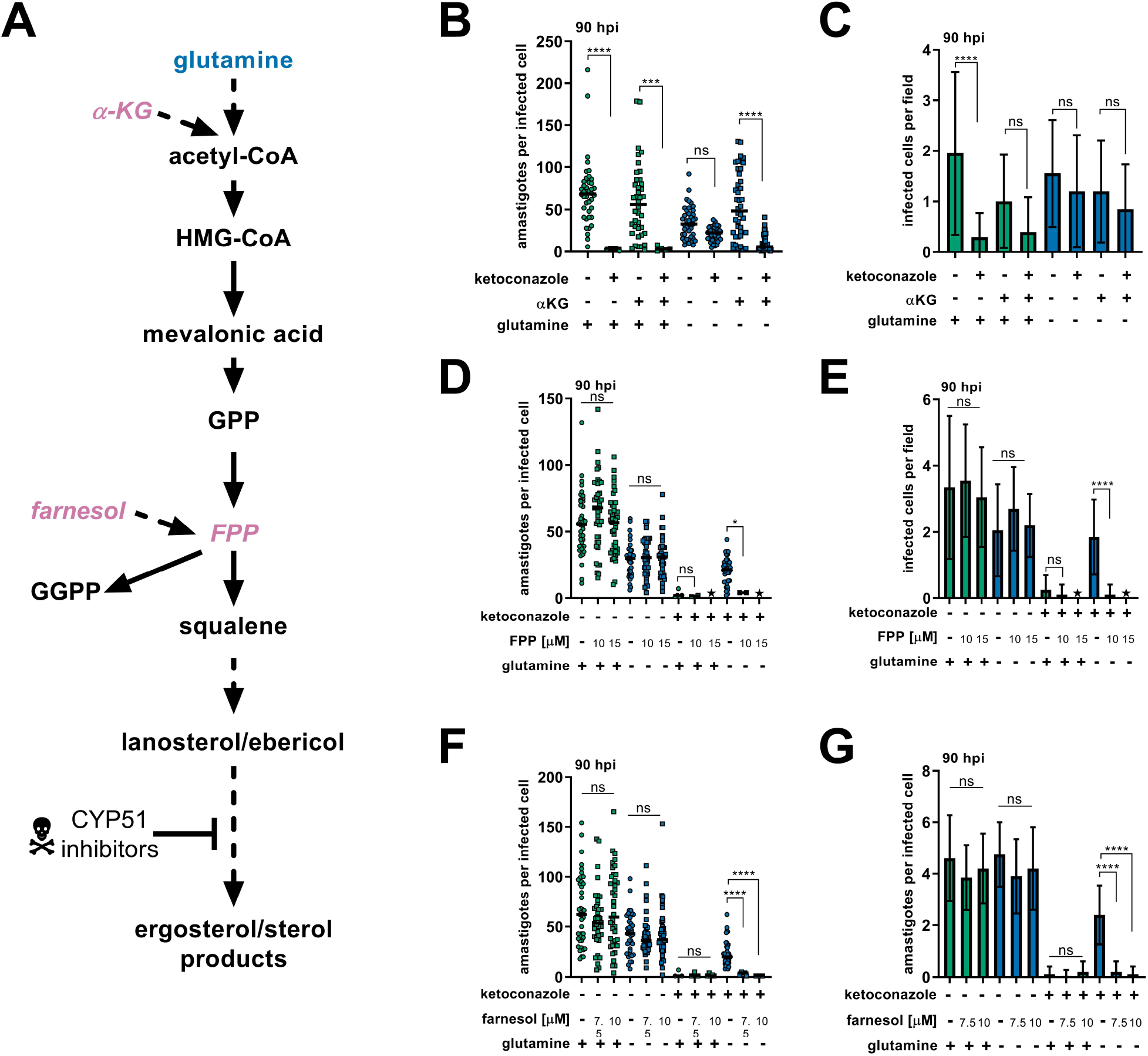
Addition of metabolites can re-sensitize intracellular *T. cruzi* amastigotes to ketoconazole in the absence of glutamine. **(A)** Schematic of endogenous sterol synthesis. Dash lined arrows indicate omission of steps for simplicity. **(B)** Microscopic counts of amastigotes per infected cell (n=40) and **(C)** infected cells per field (n=20) at 90 hpi with αKG (10 mM) supplemented where indicated. Microscopic counts of **(D)** amastigotes per infected cell (n=40) and **(E)** infected cells per field (n=20) at 90 hpi with FPP supplemented where indicated. Stars indicated conditions where parasites could not be found. **(F)** Microscopic counts of amastigotes per infected cell (n=40) and **(G)** infected cells per field (n=20) at 90 hpi with farnesol supplemented where indicated. Statistical comparisons between medians (B,D,F) were performed using a Kruskal-Wallis test with Dunn’s multiple comparisons test (****p<0.0001, ***p<0.001, *p<0.05, ns=not significant). Comparisons of means (C,E,G) were performed using a one-way ANOVA and Bonferroni’s multiple comparisons test (****p<0.0001, ns=not significant).

### *T. cruzi* glutamate dehydrogenase mutants succumb to ketoconazole in the presence and absence of supplemental glutamine

Manipulation of the growth medium, such as glutamine withdrawal or addition of metabolites, is expected to impact host cell metabolism and either directly and/or indirectly impact parasite metabolism. To assess the contribution of *T. cruzi* glutamine metabolism in sensitizing intracellular amastigotes to CYP51-targeting azoles, we took a genetic approach to dysregulate glutamine metabolism in the parasite. Using a CRISPR/Cas9-mediated approach ((Lander et al., 2015); Figure 5—figure supplement 1) we disrupted the *T. cruzi* gene encoding mitochondrial glutamate dehydrogenase (mGDH; TcCLB.509445.39), which is predicted to function in anaplerosis from glutamine (Cazzulo et al., 1979) and cytosolic (Leroux et al., 2011) isocitrate dehydrogenase (cIDH; TcCLB.506925.319) that in other systems can drive carbons from glutamine into the synthesis of sterols 22101433. Consistent with a role for mGDH in anaplerosis and fueling the TCA cycle and respiratory chain, mGDH-deficient *T. cruzi* isolated amastigotes fail to maintain ATP levels when glutamine is supplied as the sole carbon source, whereas glucose was able to sustain these mutants in a manner similar to that observed in WT parasites (Figure 5A). As compared to WT amastigotes the mGDH-deficient parasites exhibit a growth defect in mammalian cells when cultured in complete medium, but grow similar to WT amastigotes when cultured in medium without glutamine (Figure 5B). In striking contrast to WT and cIDH-deficient parasites, mGDH-deficient amastigotes fail to survive exposure to ketoconazole upon glutamine withdrawal (Figure 5B,C) with few infected cells remaining following ketoconazole treatment independent of exogenous glutamine status (Figure 5B,C). Unlike WT amastigotes, mGDH-deficient amastigotes are less sensitive to exogenous glutamine in the absence of ketoconazole (Figure 5D). In the presence of ketoconazole mGDH-deficient amastigotes show no relationship between survival and the amount of exogenous glutamine (Figure 5D). This loss of protection from ketoconazole exhibited by mGDH-disrupted *T. cruzi* amastigotes illustrates how dysregulation of parasite glutamine metabolism can influence the susceptibility of these parasites to azoles. Taken together our data show that the ability of azoles to kill intracellular *T. cruzi* amastigotes can be modulated by altering the extracellular environment or targeted inhibition of parasite metabolism, which may have broader implications for the screening and efficacy of candidate anti-trypanosomals.

**Figure 5:**
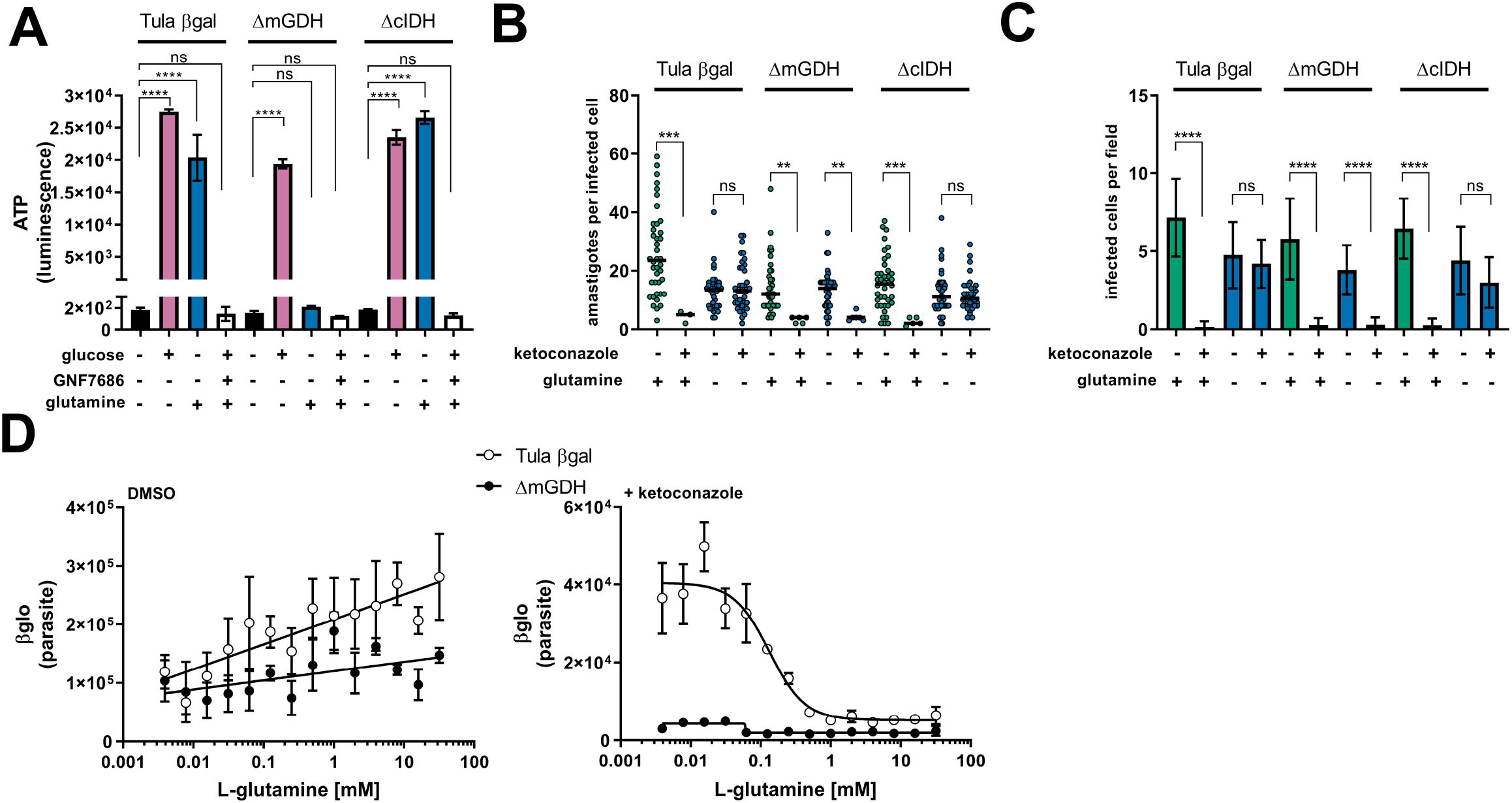
Amastigote glutamine metabolism alters sensitivity to ketoconazole. **(A)** Quantification of the levels of ATP from isolated amastigotes in carbon free buffer supplemented with glucose or glutamine. Comparisons of means, to supplemented conditions within a given parasite line, were performed using a one-way ANOVA and Bonferroni’s multiple comparisons test (p<0.0001=****, ns=not significant). **(B)** Microscopic counts of amastigotes per infected host cell (n=40) and **(C)** the number of infected cells per field (n=20) at 66 hpi. Statistical comparisons between medians (B) were performed using a Kruskal-Wallis test with Dunn’s multiple comparisons test (**p<0.01, ***p<0.001, ns=not significant). Comparisons of means (C) were performed using a one-way ANOVA and Bonferroni’s multiple comparisons test (****p<0.0001, ns=not significant). **(D)** Dose response curves of glutamine in the presence of DMSO or 5 nM ketoconazole. Mean and standard deviation shown (n=3). Tula-βgal is shown in unfilled circles and Tula-βgalΔmGDH is shown with filled circles.

## Discussion

Independent of growth rate, the metabolic state of a microorganism can be influenced by its immediate environment and have an impact of the efficacy of antimicrobials (Lopatkin et al., 2019). For pathogenic microbes this environment is largely dependent on the status of its host. Within a host, nutrient utilization and availability vary widely across tissues (Shlomi et al., 2008). Even within a single tissue, the presence of an inflammatory response shifts the local metabolism (Kominsky et al., 2010) and in many cases leads to intracellular nutrient restriction to control pathogen growth (Grohmann et al., 2017). Pathogen growth *in vitro* cannot always reflect the complete spectrum of metabolic environments present *in vivo* and consequently can confound interpretations of standard antimicrobial assays (Hicks et al., 2018; Pethe et al., 2010). These considerations are especially pertinent with regard to *T. cruzi*, a parasite that in its mammalian host replicates intracellularly in diverse tissues and persists for the lifetime of the host, exposing the parasite to an immune response that suppresses parasitemia without sterile cure (Lewis et al., 2015).

Recent clinical trials investigating the efficacy of azoles (CYP51 inhibitors) to eliminate *T. cruzi* parasitemia resulted in an initial elimination of peripheral parasitemia that unlike benznidazole was not maintained after cessation of therapy (Molina et al., 2014; Morillo et al., 2017; Torrico et al., 2018). The anti-parasitic activity of azoles without sterile cure suggests the possibility that heterogeneous environments and/or distinct populations of parasites within a single host may underlie treatment failure. We examined the influence of metabolic environment on the *in vitro* potency of benznidazole, the current first line therapy, and azoles. Benznidazole remained equally potent in all conditions tested, showing that its activity does not interact with either the slowed growth of amastigotes in these conditions or the specific nutrients tested. Unlike benznidazole, the activity of azoles was found to be affected by the absence of glutamine, but not glucose, in the medium. Rather than a shift in traditionally measured IC_50_, this change is characterized by the inability of azoles to cause radical growth reduction at all concentrations tested, potentially through drug resistance, tolerance or kill kinetics. However, this protection is not explained by increased time to kill since increasing the time of exposure to ketoconazole did not result in amastigote death in the absence of glutamine. Rather we found that intracellular amastigotes continued to proliferate in the presence of ketoconazole when glutamine is restricted and are competent to complete the lytic cycle by undergoing differentiation to trypomastigotes that egress from the host cell.

With regard to growth rate itself influencing azole efficacy, we found that slowing amastigote growth, using an inhibitor (GNF7686) of parasite cytochrome b was able to delay but not prevent amastigote death due to azoles in the absence of supplemental glutamine. In a similar scenario, more slowly growing *T. cruzi* isolates appear less susceptible to azoles in a single time point growth inhibition assay, yet more rapidly dividing strains can still outgrow following treatment (MacLean et al., 2018). An alternative explanation for parasite persistence in the presence of azoles is a complete cessation of amastigote division. While the nature of dormancy in *T. cruzi* remains under investigation (Sánchez-Valdéz et al., 2018), we report here a protective mechanism that allows for amastigote proliferation in the presence of drug at the population level. Since, slowed growth appears to induce tolerance to azoles but is insufficient to provide resistance; we investigated potential mechanisms to explain the protection from azoles mediated specifically by glutamine restriction.

Glutamine is the most abundant amino acid in the human body and has a wide intracellular distribution between tissues (Cruzat et al., 2018). Standard *in vitro* growth media compositions contain supraphysiologic amounts of glutamine to allow for the sustained growth of rapidly dividing cells. We found that *in vitro,* amastigotes do not become entirely sensitized to ketoconazole until glutamine is supplemented to levels higher than those found in plasma. Additionally, during inflammation in general, intracellular glutamine pools can become depleted in certain tissues as release rates exceed synthesis (Karinch et al., 2001). Data from this study show that *in vitro* growth conditions may belie the variable efficacy of candidate anti-parasitic compounds and offer complementary approaches to better prioritize new candidates.

Our data and others show that azoles act only after several rounds of parasite division, likely through the gradual depletion of sterol end products and/or the buildup of 14-methylated sterol synthesis intermediates. Similar to fungal species, *T. cruzi* endogenously synthesized ergostane-type sterols. The fungal/parasite cytochrome P450 14α-demethylase (CYP51/Erg11) is involved in the synthesis of ergosterol and is the target of azole drugs. Drug resistance to azoles observed in fungal pathogenesis include mutations in CYP51 (Howard et al., 2009), drug efflux (Prasad and Rawal, 2014), selection for sterol auxotrophy (Hazen et al., 2005) or suppressor mutations that alter the composition of 14-methylated sterol synthesis intermediates (Kelly et al., 1995). Target site or suppressor mutations cannot explain protection mediated by glutamine restriction in *T. cruzi* amastigotes because we found that amastigotes are protected as a population, which occurs rapidly within a single lytic cycle. Under these conditions, amastigotes are not exposed to prolonged selection. Protection from azoles mediated by glutamine withdrawal is likely not due to a decrease in activity due to drug efflux, since the generation of sterols downstream of CYP51 is abolished in the absence of glutamine, demonstrating that the activity of ketoconazole is unchanged. These data demonstrate that protection may not be mediated by changes to the activity or sensitivity of CYP51 to azoles but rather changes to the consequences of CYP51 inhibition.

Increased membrane fluidity and heat sensitivity seen in *Leishmania major* CYP51 knockouts (Xu et al., 2014) but not in knockouts of sterol methyltransferase (Mukherjee et al., 2019) suggests that accumulation of 14-methylated sterols rather than the absence in ergosterol effects parasite viability. We found that the carbons from glutamine enter the endogenous sterol synthesis pathway in *T. cruzi* amastigotes and are incorporated into the 14-methylated sterol synthesis intermediates lanosterol and ebericol. The incorporation of these carbons into amastigote sterols suggests that removal of glutamine has the potential to diminish flux through the sterol synthesis pathway. While lanosterol and ebericol both increase in the presence of ketoconazole, the relative amount of lanosterol is less when amastigotes are grown in the absence of glutamine. Since we have only measured free sterols in isolated *T. cruzi* amastigotes, it is possible that sterols or their synthesis intermediates are esterified (Pereira et al., 2018; Taylor and Parks, 1978) or exported from the amastigote and therefore not detected using these methods. In line with glutamine modulating flux through the sterol biosynthesis pathway, addition of metabolites upstream of CYP51 (α-KG, FPP, farnesol) were shown to re-sensitize amastigotes to the cytotoxic effects of ketoconazole in the absence of glutamine. Taken together these data show that carbons derived from glutamine may determine flux through the endogenous sterol synthesis pathway in amastigotes and influence the buildup of 14-methylated species.

Within the context of our proposed model, our finding that mGDH-deficient *T. cruzi* amastigotes fail to survive exposure to ketoconazole upon glutamine restriction suggests that mutant parasites are unable to reduce flux of carbons into the sterol synthesis pathway in the absence of glutamine. Although the precise contribution of parasite and host metabolism in modulating sterol biosynthesis in intracellular amastigotes is still undetermined, our data clearly indicate that parasite glutamine metabolism plays a critical role in sensitizing *T. cruzi* to azoles.

In summary, we find that the sensitivity of intracellular amastigotes to azoles is modulated by their glutamine metabolism, characterized by changes in flux through the sterol synthesis pathway. These observations have implications for *T. cruzi* antimicrobial prioritization and further evidence that the metabolic state of a microorganism is an important consideration for determining drug susceptibility. Even though the identification of new targets for antiparasitic compounds (Khare et al., 2016, 2015b) is promising, a better understanding of parasite metabolism and reasons for failure of prior candidates has the potential to aid in the prioritization of these potential therapies. In addition, the ability to modulate drug susceptibility through nutrient availability *in vitro* suggests that nutrient supplementation *in vivo* should be explored as a potential combination therapy.

## Materials and Methods

### Mammalian cell culture

Mammalian cells were maintained at 37°C in a 5% CO_2_ incubator. Dulbecco’s modified Eagle medium (DMEM; HyClone, Logan, Utah) supplemented with 10% FBS (Gibco, Waltham, Massachusetts), 25 mM glucose, 2 mM L-glutamine and 100U/mL penicillin-streptomycin was used for propagated for uninfected cultures (DMEM-10). Unless stated otherwise, cultures infected with *Trypanosoma cruzi* were maintained in DMEM with 2% FBS (DMEM-2). Normal Human Neonatal Dermal Fibroblasts (NHDF; Lonza, Basel, Switzerland) were passaged prior to reaching confluence.

### Parasite maintenance

Tulahuén LacZ clone C4 (Tula-βgal), PRA-330 (ATCC, Manassas, Virginia) was obtained directly and passaged weekly in LLC-MK_2_, CCL-7 (ATCC, Manassas, Virginia) cells (Buckner et al., 1996). Trypomastigotes were prepared by collecting the supernatant from infected cultures and centrifuging for 10 min at 2,060 x g followed by incubation at 37°C for >2 hours to allow for trypomastigotes to swim from the pellet. After incubation the supernatant containing trypomastigotes was collected and washed in DMEM-2, enumerated using a Neubauer chamber and used for subsequent infections.

### Quantification of parasite load by luminescence

Tula-βgal parasite load was measured using luminescence as described previously (Caradonna et al., 2013; Shah-Simpson et al., 2017). One day prior to infection NHDFs were seeded in 384-well plates (Corning, Corning, New York) at a density of 1,500 cells per well and allowed to attach. Purified trypomastigotes were added at a multiplicity of infection (MOI) of 1.25 and allowed to invade for 2 hours, followed by two washes with PBS and subsequent addition of DMEM-2 without phenol red. Treatments were initiated at 18 hours post infection (hpi) to avoid any potential impacts of trypomastigote invasion and/or differentiation. At the indicated time points growth media was removed and 10 μl Beta-Glo (Promega, Madison, Wisconsin) was added per well. Plates were incubated for >30 min at room-temperature to allow the reaction to reach equilibrium and read using an EnVision plate reader (PerkinElmer, Waltham, Massachusetts). Luminescence from uninfected wells was determined for each treatment and subtracted from infected wells to account for signal not derived from parasites.

### Compound and supplement stocks

Compounds were purchased and diluted to stock concentrations: Ketoconazole (Enzo, Farmingdale, New York) 15 mM stock in DMSO, Ravuconazole (Sigma, St. Louis, Missouri) 15 mM DMSO, Itraconazole (BioVision, Milpitas, California) 15 mM DMSO, GNF7686 (Vitas-M Laboratory, Champaign, Illinois) 5 mM stock in DMSO, FPP (Sigma, St. Louis, Missouri) 2.3 mM stock in methanol, Farnesol (Sigma, St. Louis, Missouri) 100 mM in ethanol, NAC (Sigma, St. Louis, Missouri) 200 mM in DMEM base, Glutathione (Sigma, St. Louis, Missouri) 162 mM in media.

### Microscopy

Host cells were seeded one day prior to infection on coverslips (EMS, Hatfield, Pennsylvania) in 24-well plates at a density of 4 × 10^4^ cells per well. Cells were infected for 2 hours at a MOI of 2 and subsequently washed twice with PBS followed by addition of DMEM-2. Coverslips were fixed in 1% PFA-PBS and stained in a 0.1% Triton X-100–PBS solution containing 100 ng/ml DAPI (Sigma, St. Louis, Missouri) for 5 min. After staining coverslips were washed with PBS and mounted with ProLong Antifade (Thermo Fisher, Waltham, Massachusetts) on glass slides. Amastigotes were counted using a Nikon eclipse TE300. Amastigotes per infected host cell and the number of infected host cells per microscopic field were recorded.

### Western Blot

Uninfected cells were lysed in 1 mL M-PER Mammalian Protein Extraction Reagent (Thermo Fisher, Waltham, Massachusetts) directly in culture wells and boiled for 10 minutes. Soluble lysate (50 μg) was loaded onto a 10% Mini-Protean TGX Gel (Bio-Rad, Hercules, CA). Proteins were transferred to a nitrocellulose membrane and blocked with a 1:1 dilution of SEA BLOCK (Thermo Fisher, Waltham, Massachusetts):PBS overnight at 4°C. The membrane was probed in blocking buffer with anti-Hif1a EPR16897 (1:1,500) (Abcam, Cambridge, MA) and anti-βactin (Sigma, St. Louis, Missouri) (1:1,000) for 1h at room temperature in hybridization tubes. After probing the membrane was washed in 1X PBS for 30 minutes, replacing PBS every 5 minutes for a total of 6 washes. Secondary antibodies, anti-mouse DyLight 680 (Cell Signaling, Dancers, MA) (1:15,000) and anti-rabbit Dylight 800 (Thermo Fisher, Waltham, Massachusetts) (1:10,000) were added and incubated for 1 hour at room temperature. The membrane was visualized using a LI-COR imaging system (LI-COR, Lincoln, NE).

### Gene Disruption

Gene disruption was accomplished using a homology directed repair, Cas9 mediated system modified from (Lander et al., 2015). The Cas9/gRNA expression construct pTREX-n-Cas9 was first modified using the Q5 mutagenesis kit (NEB, Ipswich, Massachusetts), primers 5’-CCCAAAAAGAAAAGGAAGGTTGATTAGAAGCTTATCGATACCGTCGAC-3’ and 5’-GTCCTCGACTTTTCGCTTCTTTTTCGGGTCGCCTCCCAGCTGAGA-3’, to remove the GFP and HA tags from Cas9 and maintain the SV40-NLS. Guide RNA design used the EuPaGDT system (Peng and Tarleton, 2015). A guide RNA targeting sequence specific to GFP and scaffolding were amplified by PCR, 5’-TATAGGATCCCAGATTGTGTGGACAGGTAA-3’ and 5’-CAGTGGATCCAAAAAAGCACCGACTCGGTG-3’, using pUC_sgRNA as a template as described in (Lander et al., 2015) and cloned into pTREX-n-Cas9-noTags using BamHI. Mutations to the specificity of guides in pTREX-n-Cas9-noTags was performed using a Q5 mutagenesis kit (NEB, Ipswich, Massachusetts). Guides targeting the mitochondrial glutamate dehydrogenase (mGDH, TcCLB.509445.39) were inserted using 5’-TGTTACACGGGTTTTAGAGCTAGAAATAGC-3’/5’-AAGGCCGTAGGGATCCACTAGAACTCTTG-3’ for g195 and 5’-CAAAGGCCGTGTTTTAGAGCTAGAAATAGC-3’/5’-TTACACGGAGGGATCCACTAGAACTCTTG-3’ for g216. Guides targeting the cytosolic isocitrate dehydrogenase (cIDH, TcCLB.506925.319) were inserted using 5’-GAGCCTCGTCGTTTTAGAGCTAGAAATAGC-3’/5’-GTGTAAGGGAGGATCCACTAGAACTCTTG-3’ for g110rc and 5’-GAAGCAGATGGTTTTAGAGCTAGAAATAGC-3’/5’-AAGTTGAACTGGATCCACTAGAACTCTTG-3’ for g110. Donor DNA containing drug resistant was generated by PCR using ultramers with 100bp homology to regions flanking the predicted Cas9 cut site and an in frame P2A ribosomal skip peptide.

Epimastigotes were grown at 27°C in liver infusion tryptose (LIT) containing 10% FBS. Log-phase epimastigotes were transfected simultaneously with pTREX-Cas9-gRNA plasmids and donor DNA using an Amaxa nucleofector, program U-33 in Tb-BSF (Schumann Burkard et al., 2011). Transfected epimastigotes were allowed to recover for 24 hours and subsequently cloned in the presence of drug corresponding to the resistance marker present in the donor DNA. Clones were screening using primers, 5’-ATGCGGCGTGTGGTTATTATGG-3’/5’-TGGCTCCTTTAAAAGAAGCGCG-3’ (mGDH) and 5’-CGTGACAAAACGGACGATCAGG-3’/5’-TTCTGATCCAGTTTGCCCCGAT-3’ (cIDH) that flank the insertion site.

### Sterol Extraction

The method for extraction of sterols was based on protocols described in (Sharma et al., 2017). Extraction occurred in glass PYREX tubes (Corning, Corning, New York) and all solvents used were HPLC grade or higher. Lipids were first extracted three times from cell pellets using C:M (2:1, v/v) and centrifuged each time at 1,800 x g for 15 min at 4°C followed by collection of the supernatant in new tubes. The supernatant was dried under a constant stream of N_2_ and the resulting material was subjected to a Folch’s partitioning (4:2:1.5, C:M:W). The lower phase was removed, dried under N_2_ and re-suspended in chloroform, passed over a silica 60 column and eluted with chloroform.

### Radiolabeling and thin layer chromatography

For radiolabeling 25μCi/condition of universally labeled ^14^C-glutamine (Moravek, Brea, California) with a specific activity of 281 mCi/mmol was added to DMEM-2 with 0.5 mM unlabeled glutamine. Infection, amastigote isolation and sterol extraction were carried out as described. Separation of species by thin layer chromatography (TLC) was accomplished through a protocol modified from (Andrade-Neto et al., 2011). Silica gel on TLC aluminum foils (Honeywell Fluka, Charlotte North Carolina) were pre-conditioned with a silver nitrate (1% w/v) methanol solution and allowed to dry. Samples and standards were added to TLC plates and first placed in a tank containing a mobile phase of hexane:ethyl ether:acetic acid (60:40:1) and solvent was allowed to reach half way up the plate. Subsequently plates were placed in a second tank with a mobile phase of hexane:chloroform:acetic acid (80:20:1) until the mobile phase reached near the top of the plate. Plates were exposed for seven days to phosphorimager screens and imaged using a Typhoon Phosphorimager-FLA7000 (GE Healthcare, Chicago, Illinois). Prior to charring at 100°C for 2-5 min plates were soaked in a ferric chloride water solution (50mg FeCl_3_/100mL) with 5% acetic acid and 5% sulfuric acid (Lowry, 1968).

### GC-MS

GC/MS analysis was performed on a Thermo Scientific TRACE 1310 Gas Chromatograph equipped with a Thermo Scientific Q Exactive Orbitrap mass spectrometry system. 50μL of the (BSTFA+10% TMCS)/pyridine (5/1 v/v) was added into each vial, vortexed well, and heated at 70°C for 30 min. 1 μL sample was injected into a Thermo fused-silica capillary column of cross-linked TG-5SILMS (30 m × 0.25 mm × 0.25 μm). The GC conditions were as follows: inlet and transfer line temperatures, 290°C; oven temperature program, 50°C for 0 min, 24°C/min to 325°C for 5.7 min; inlet helium carrier gas flow rate, 1 mL/min; split ratio, 5. The electron impact (EI)-MS conditions were as follows: ion source temperature, 310°C; full scan m/z range, 30 - 750 Da; resolution, 60,000; AGC target, 1e6; maximum IT, 200ms. Data were acquired and analyzed with Thermo TraceFinder 4.1 software package. Standards for cholesterol, ergosterol, lanosterol, episterol and zymosterol were used for identification. Universal ^13^C-glutamine was re-suspended to a stock concentration of 200 mM in water (Cambridge Isotope Labs, Tewksbury, Massachusetts). Prior to sterol extraction sitosterol-d7 (Avanti Polar Lipids, Alabaster, Alabama) was add as an internal standard (ISTD) at 1.12 μg/2e7 isolated amastigotes.

### Amastigote Isolation

Infected monolayers were washed 2 times with PBS and cell detachment was achieved using a sterol free dissociation reagent, Accumax (Innovative Cell Technologies, San Diego, California). Cell suspensions were washed 2 times with PBS by centrifugation at 700 x g for 10min at 4°C. The resulting cell pellets were lysed by passage through a 28-gauge needle or using the Miltenyi GentleMACS dissociator (M tubes, Protein_01 protocol). Lysate was passed over a PD-10 column (GE Healthcare, Chicago, Illinois) equilibrated with PBS. Eluted parasites were washed three times in PBS by centrifugation at 2300 x g at 4°C.

### Clonal outgrowth

Measurement of clonal outgrowth utilized a modified protocol from (Dumoulin and Burleigh, 2018) to allow for detection by luminescence. Host cells were seeded in 384 well plates and 25 trypomastigotes per well were allowed to invade for 2 hours, followed by 2 washes with PBS to removed uninvaded trypomastigotes. Treatments were initiated at 18 hpi and wells were washed at 66 hpi twice with PBS followed by addition of DMEM-2. Cultures were allowed to grow for 14 days and subsequently measured for presence of parasites by luminescence as described previously.

### ATP assay

Isolated amastigotes were re-suspended in KHB as described (Shah-Simpson et al., 2017). After isolation, 8e5 amastigotes per well were added to 96 well plates. Where indicated glucose was supplemented to a final concentration of 25 mM, glutamine at 2 mM and GNF7686 at 2.5 μM. Amastigotes were incubated at 37°C for 72 hours followed by lysis and measurement of luminescence using the ATPlite assay (Perkin Elmer, Waltham, MA).

## Supporting information

Figure 1_Suppplemental 4

## Figure Legends

**Figure 1 –figure supplement 1: Experimental schematic for *in vitro* infection and readouts**

Trypomastigotes (Tula-βgal) are incubated with mammalian host cells for 2 hours to allow invasion. Remaining extracellular parasites are subsequently removed by thorough rinsing of monolayers. Internalized parasites undergo differentiation into mature amastigotes and any treatments or media adjustments are initiated at 18 hpi prior to the first amastigote division. At indicated time points post-infection (e.g. 42-90 hpi), infected cultures have one of several fates depending on the experiment, as illustrated and described in detail in the Methods.

**Figure 1 –figure supplement 2: Sensitivity to additional azole drugs is modulated by glutamine**

**(A)** Dose response curves of itraconazole, **(B)** posaconazole, and **(C)** ravuconazole treatment measured at 66 hpi. Treatment including media compositions are indicated and growth is normalized to the largest mean in each data set. Mean (circles) and standard deviation show (n=4).

**Figure 1 –figure supplement 3: Ketoconazole does not prohibit amastigote proliferation at 42 hpi**

Microscopic counts of intracellular amastigotes per host cell at 42 hpi following treatment at 18 hpi with ketoconazole (5 nM) under the indicated conditions.

**Figure 1 –figure supplement 4: Increased exposure to ketoconazole does not change clonal outgrowth**

Detection of clonal outgrowth 14 days after the indicated treatments, normalized to DMSO treatment. A pulse length of 48 hr (18 hpi – 66 hpi) or 72 hr (18 hpi – 90hpi) does not alter sensitivity to ketoconazole in complete medium or medium without glutamine.

**Figure 1 –figure supplement 5: Proline or Histidine supplementation do not sensitize amastigotes to ketoconazole in the absence of glutamine**

Dose response curves of proline/histidine in the absence of supplemental glutamine (n=3).

**Figure 2 –figure supplement 1: Antioxidants or hypoxia do not alter sensitivity of amastigotes from ketoconazole**

**(A)** Dose response of N-acetylcysteine and **(B)** glutathione measured at 66 hpi in the indicated treatment conditions. Mean and standard deviations are show (n=2). **(C)** Microscopic counts of the number of amastigotes per infected host cell, mean indicated, (n=40) and **(D)** the number of infected cells per 20 fields, mean and standard deviation shown (n=3). Growth in complete medium under normoxia (20% atmospheric oxygen) or hypoxia (1.3% oxygen) and ketoconazole (5 nM) where indicated. **(E)** Western blot of uninfected whole host cell lysate. Hif1α is induced under hypoxia and in the presence of DMOG (0.8 mM for 6 hours) as a positive control. Statistical comparisons between medians (C) were performed using a Kruskal-Wallis test with Dunn’s multiple comparisons test (****p<0.0001, ns=not significant). Comparisons of means (D) were performed using a one-way ANOVA and Bonferroni’s multiple comparisons test (****p<0.0001, ns=not significant).

**Figure 3 –figure supplement 1: Time course establishes 52 hpi as optimal time point to harvest intracellular amastigotes following ketoconazole treatment**

Time course following treatment with ketoconazole (5 nM) in complete media. Amastigotes per infected host cell (n=40) and infected cells per 20 fields are shown. 52 hpi identified as maximum time of ketoconazole exposure prior to measurable loss of intracellular amastigotes.

**Figure 3 –figure supplement 2: Endogenous lanosterol and ebericol but not host derived cholesterol are reliable quantifiable from isolated intracellular amastigotes**

Isolated amastigotes (52 hpi) were prepared on three independent occasions (biological) and each isolation was extracted three separate times (sterol extraction). The coefficient of variation of standard normalized area (GC-MS) was determined, means indicated. Variation in cholesterol between biological replicates prohibits reliable quantification.

**Figure 4 –figure supplement 1: FPP and farnesol re-sensitize amastigotes to ketoconazole by 66 hpi**

**(A)** Microscopic counts of amastigotes per infected cell (n=40) and **(B)** infected cells per field (n=20) at 66 hpi with FPP supplemented where indicated. **(C)** Microscopic counts of amastigotes per infected cell (n=40) and **(D)** infected cells per field (n=20) at 66 hpi with farnesol supplemented where indicated. Statistical comparisons between medians (A,C) were performed using a Kruskal-Wallis test with Dunn’s multiple comparisons test (****p<0.0001, ns=not significant). Comparisons of means (B,D) were performed using a one-way ANOVA and Bonferroni’s multiple comparisons test (****p<0.0001, ns=not significant).

**Figure 5 –figure supplement 1: Verification of gene disruptions**

**(A)** Schematic for the generation of Cas9 mediated double strand breaks (black star) and recombination using homology directed repair. **(B)** PCR verification of selected clones to verify target integration.

## Acknowledgements

We wish to acknowledge the ICCB-Longwood Screening Facility at Harvard Medical School for help with optimization of plate-based luminescence assays. We also thank Dr. Igor C. Almeida and Dr. Lucas Pagura for help with sterol extraction and thin layer chromatography protocols. This work was funded by NIH NIAID R21 AI146815-01 awarded to B.A.B. and American Heart Association Founders Affiliate Postdoctoral fellowship 19POST34380209 awarded to P.C.D.

