## Supplementary figures and images for "Glutamine metabolism modulates azole susceptibility in *Trypanosoma cruzi* amastigotes"

### Figure 1_Supplemental 1

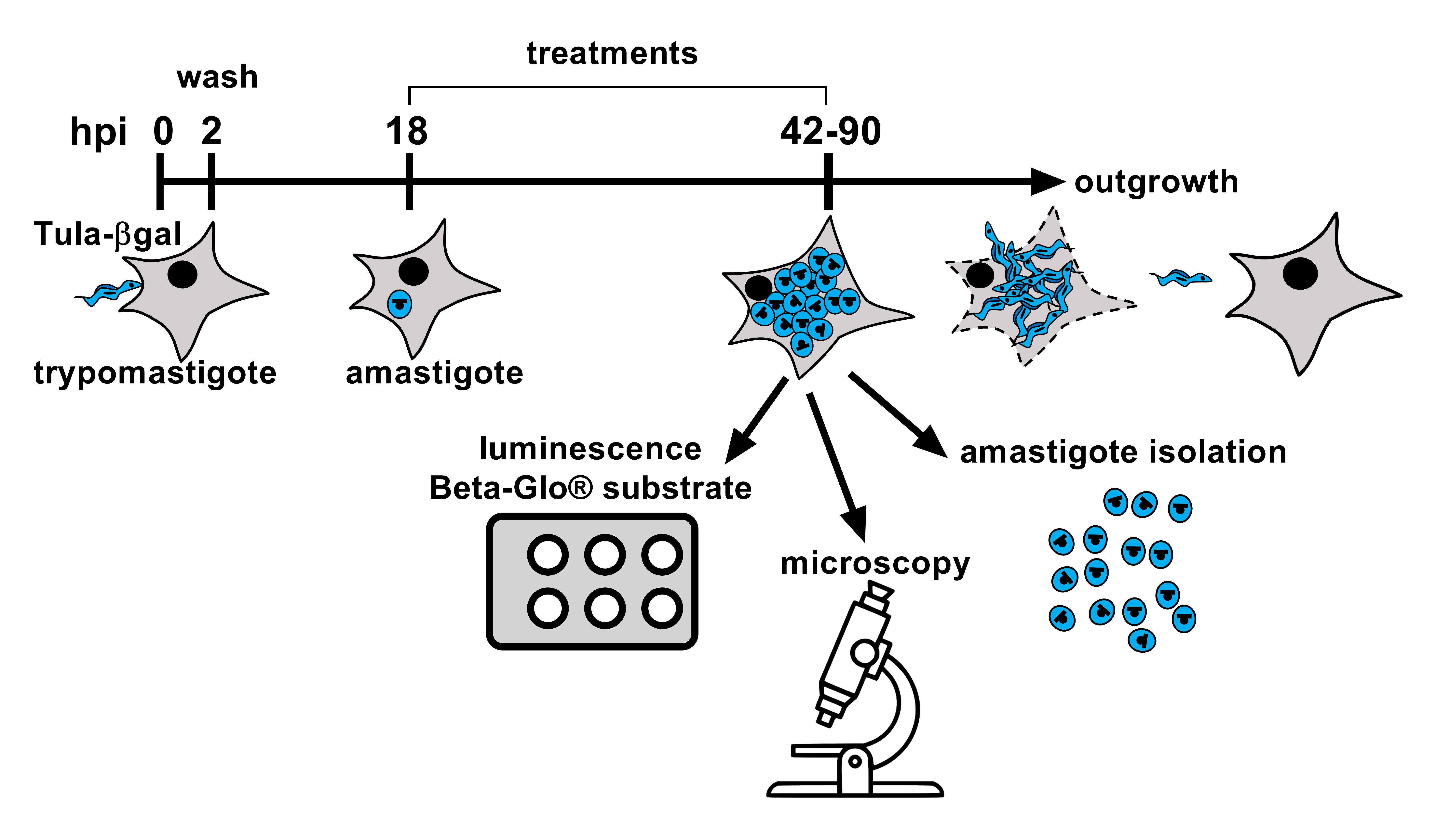

### Figure 1_Supplemental 2

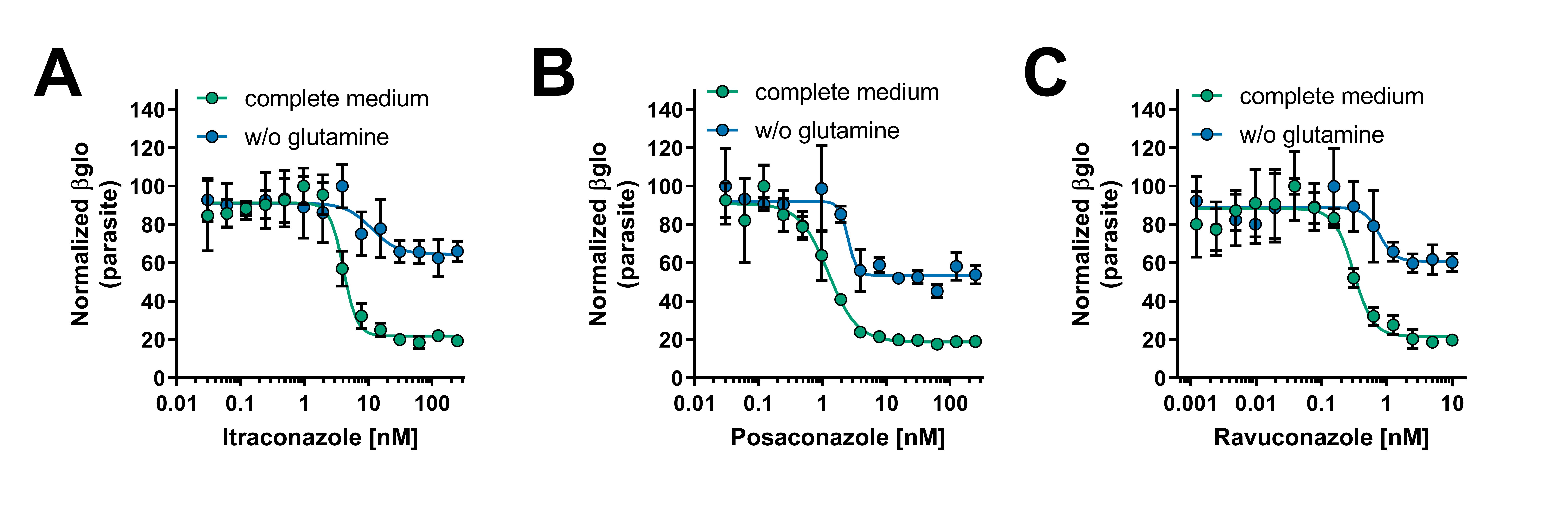

### Figure 1_Supplemental 3

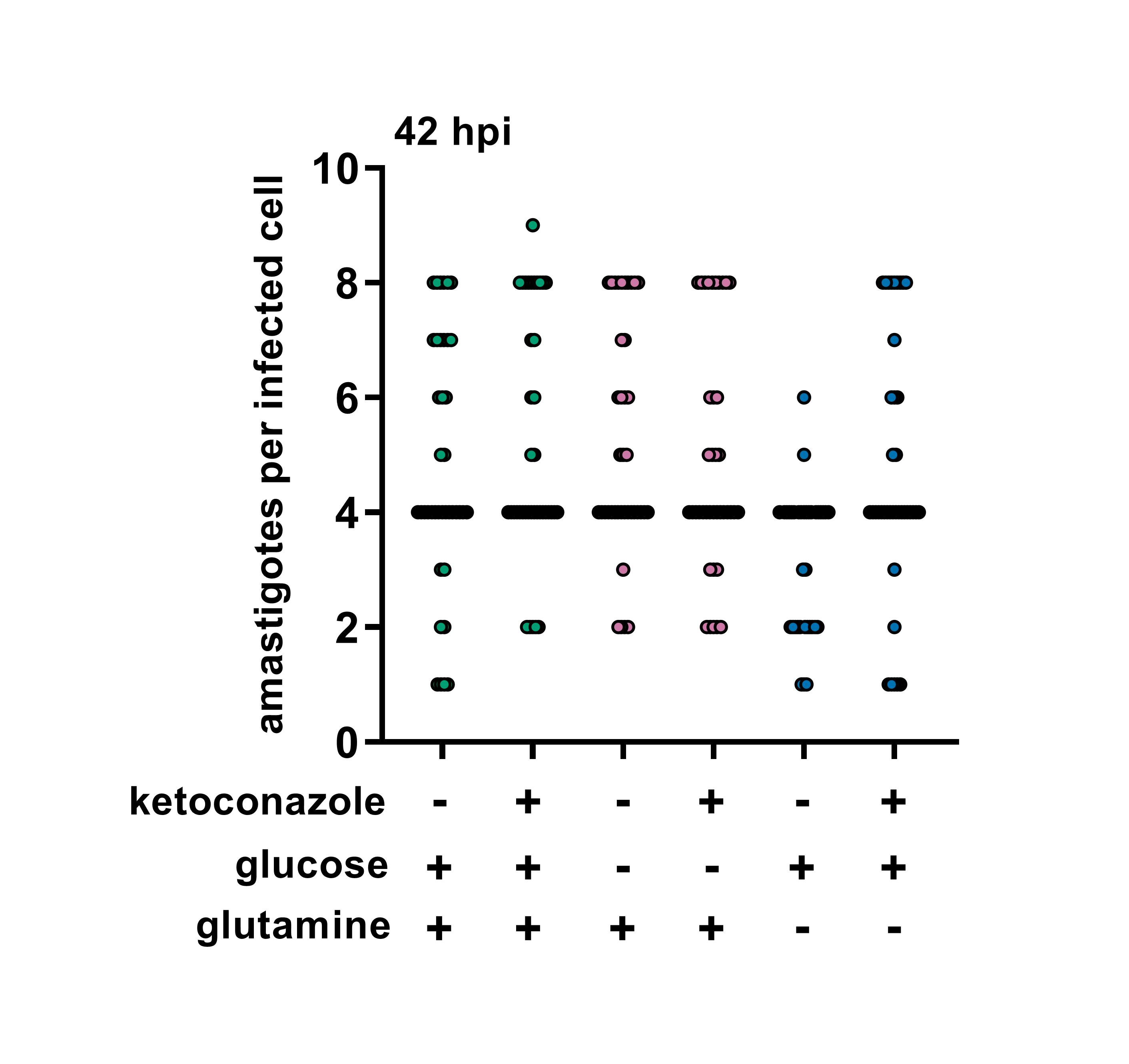

### Figure 1_Supplemental 5

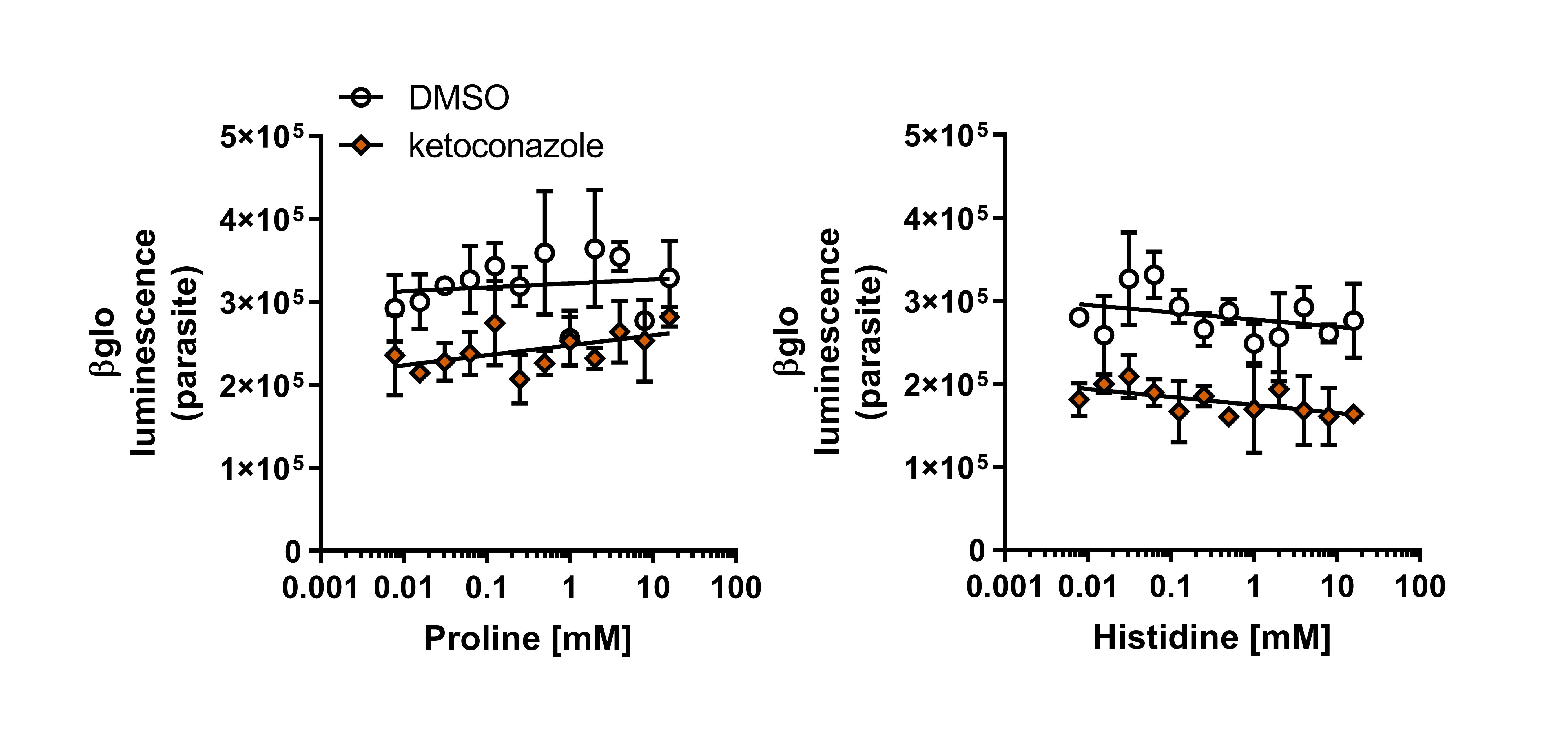

### Figure 1_Suppplemental 4

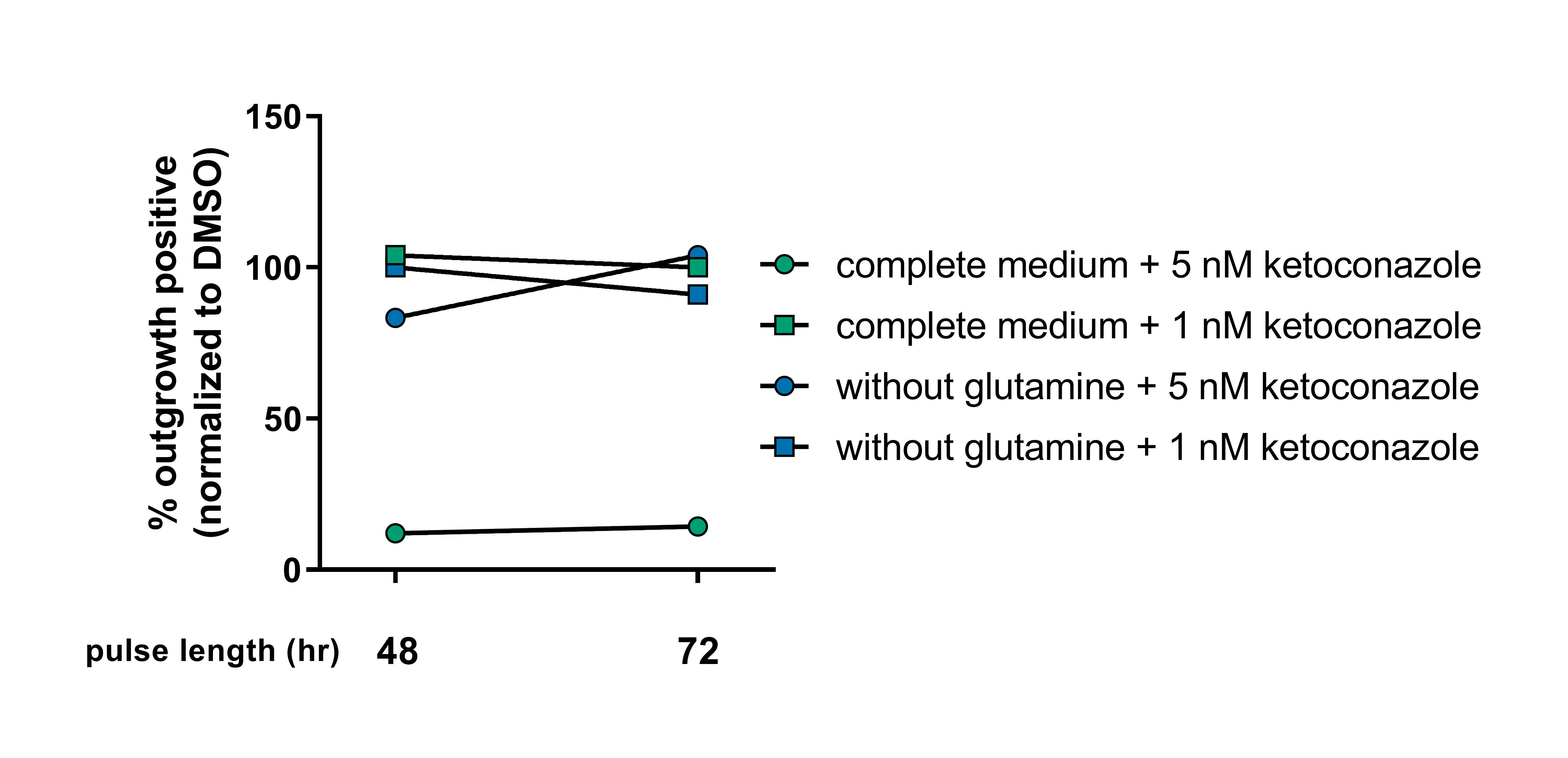

### Figure 2_Supplemental 1

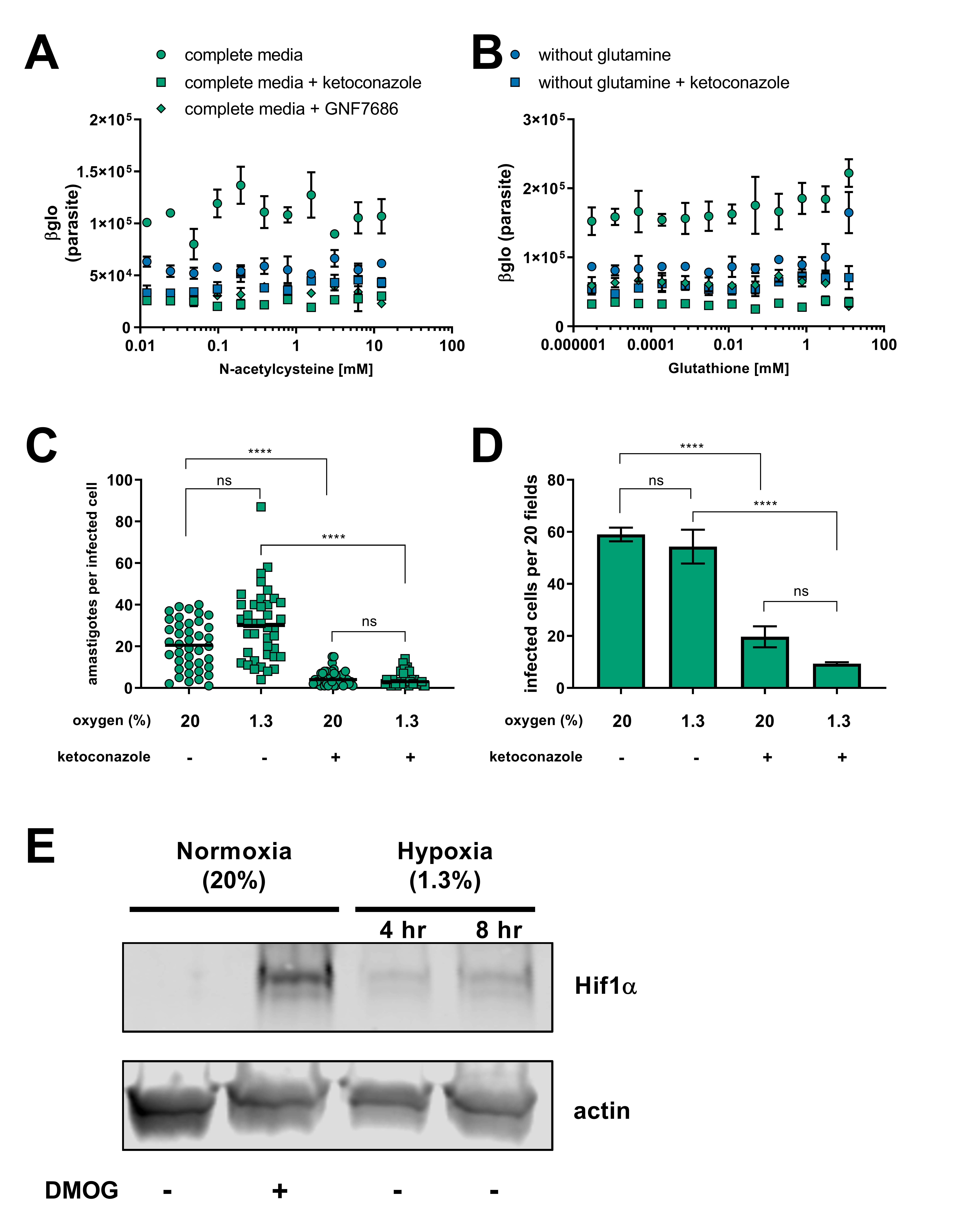

### Figure 3_Supplemental 1

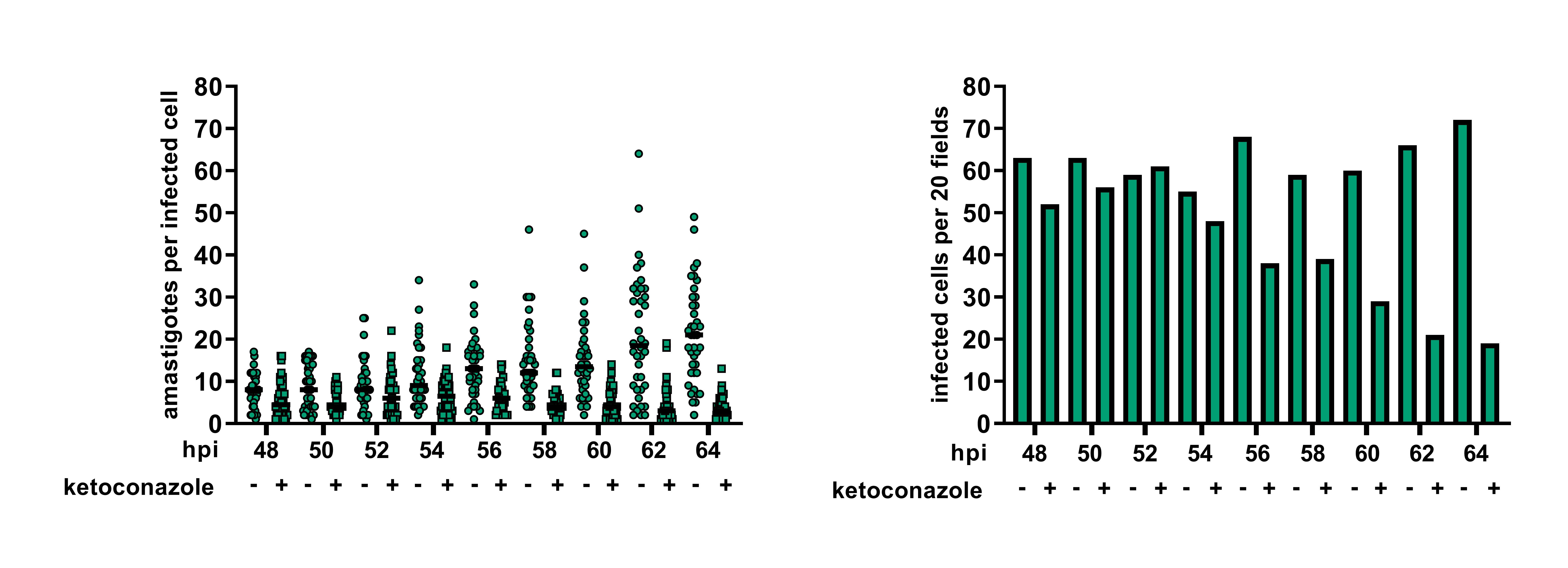

### Figure 3_Supplemental 2

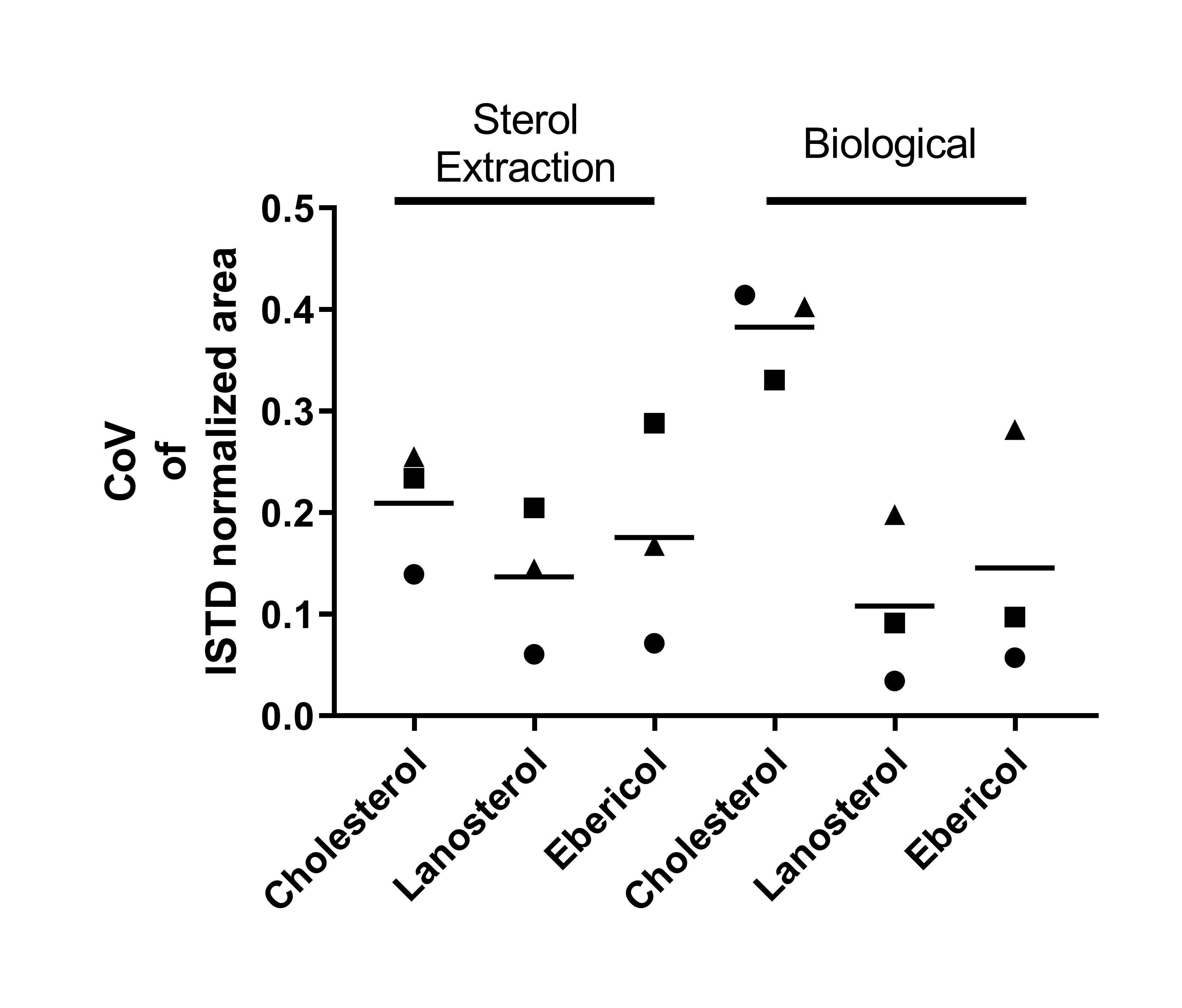

### Figure 4_Supplemental 1

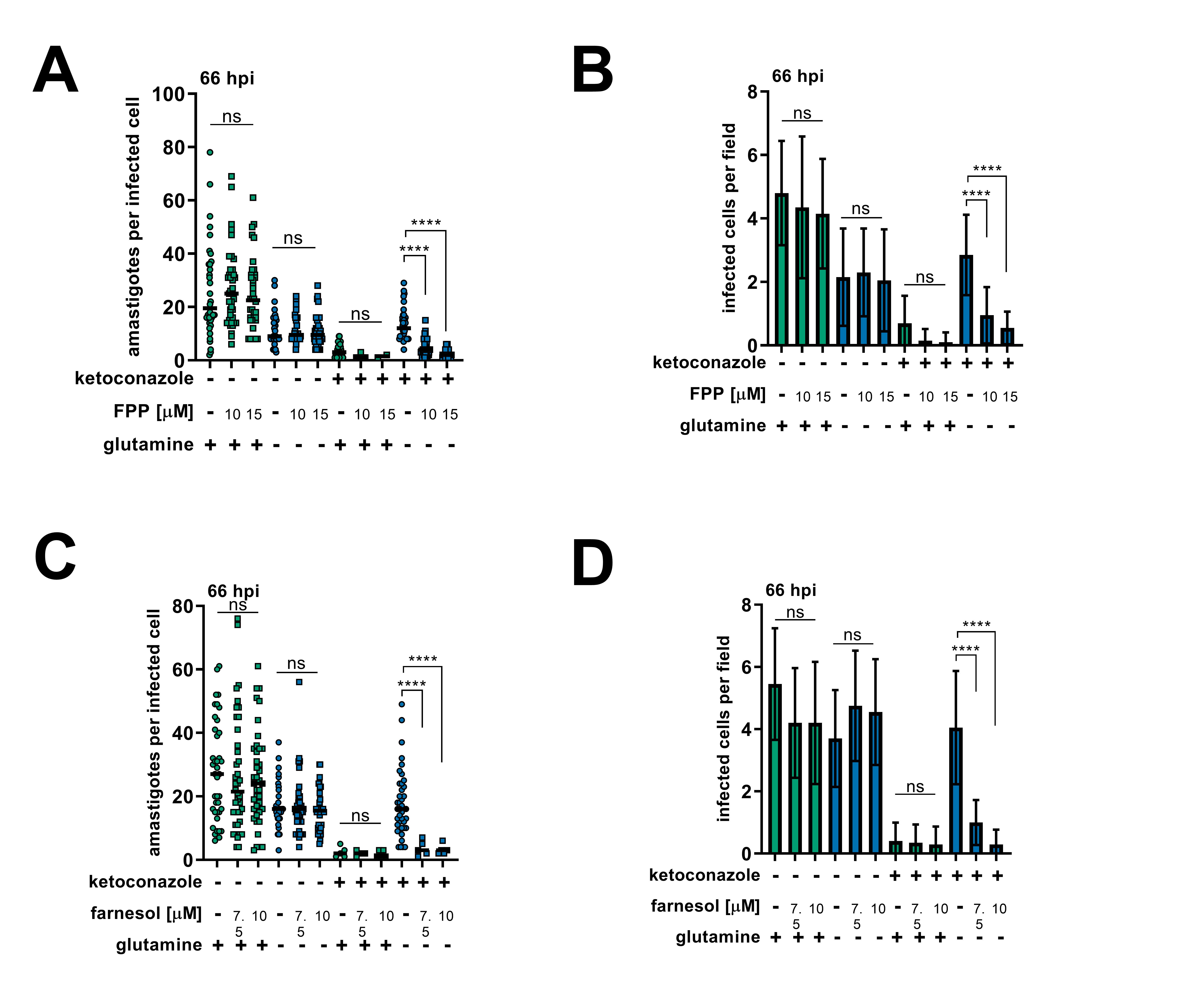

### Figure 5_Supplemental 1

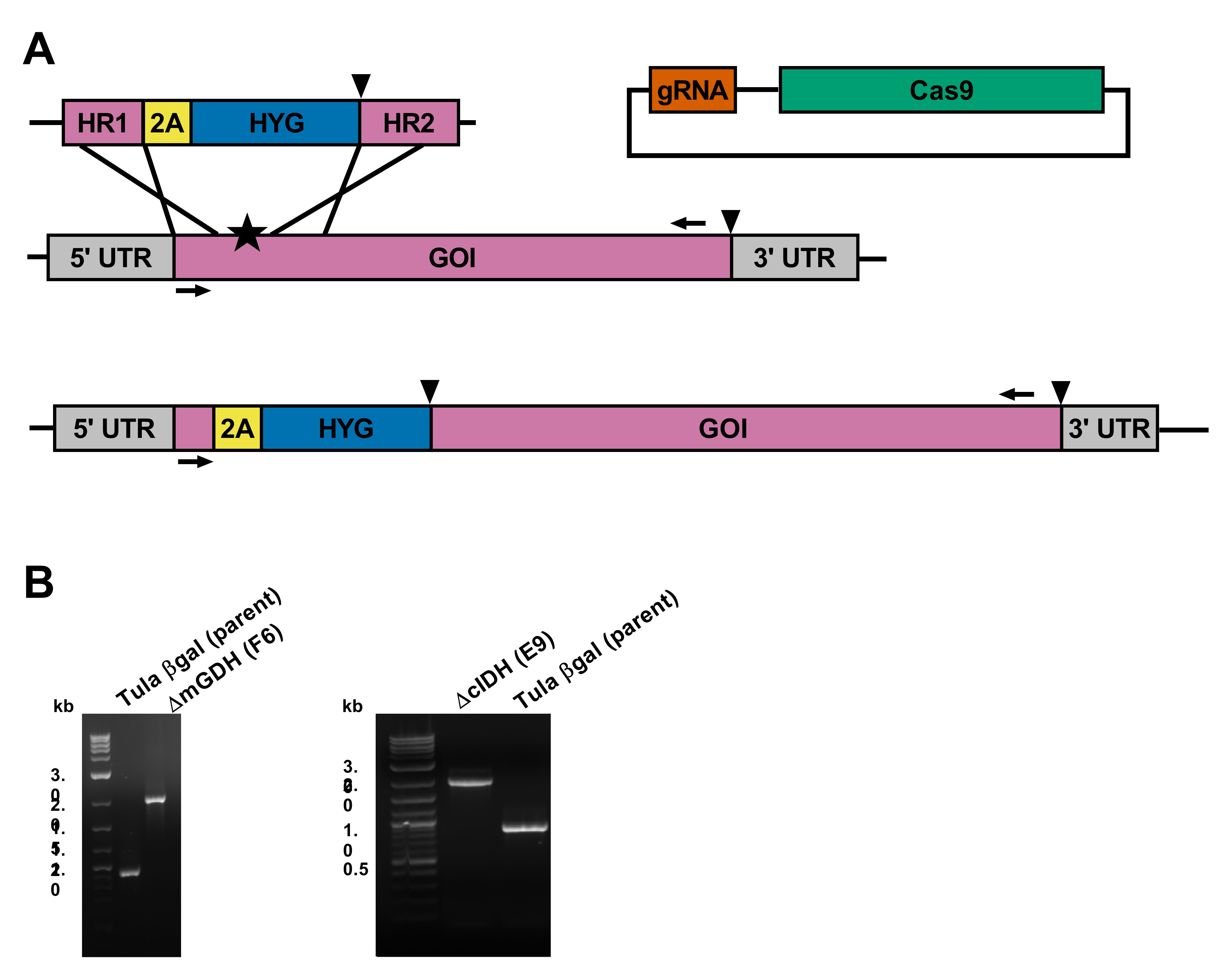
